# DyNDG: Identifying Leukemia-Related Genes based on the Time-Series Dynamical Network by Integrating Differential Genes

**DOI:** 10.1101/2024.11.24.624224

**Authors:** A Jin, Ju Xiang, Xiangmao Meng, Yue Sheng, Hongling Peng, Min Li

**Affiliations:** School of Computer Science and Engineering, Central South University, Changsha 410083, China; School of Computer and Communication Engineering, Changsha University of Science & Technolog, Changsha 410114, China; School of Computer Science & School of Cyberspace Science, Xiangtan University, Xiangtan, 411105, China; Department of Hematology, The Second Xiangya Hospital of Central South University, Changsha, 410011, China; Hunan Engineering Research Center of Cell Immunotherapy for Hematopoietic Malignancies, Changsha 410011, China

**Keywords:** Leukemia, Dynamic network, Random walk, Differential expressed genes

## Abstract

Leukemia is a malignant disease of progressive accumulation characterized by high morbidity and mortality rates, and investigating its disease genes is crucial for understanding its etiology and pathogenesis. Network propagation methods have emerged and been widely employed in disease gene prediction, but most of them focus on static biological networks, which hinders their applicability and effectiveness in the study of progressive diseases. Moreover, there is currently a lack of special algorithms for the identification of leukemia disease genes. Here, we proposed DyNDG, a novel Dynamic Network-based model, which integrates Differentially Expressed Genes to identify leukemia-related genes. Initially, we constructed a time-series dynamic network to model the development trajectory of leukemia. Then, we built a background-temporal multilayer network by integrating both the dynamic network and the static background network, which was initialized with differentially expressed genes at each stage. To quantify the associations between genes and leukemia, we extended a random walk process to the background-temporal multilayer network. The experimental results demonstrate that DyNDG achieves superior accuracy compared to several state-of-the-art methods. Moreover, after excluding housekeeping genes, DyNDG yields a set of promising candidate genes associated with leukemia progression or potential biomarkers, indicating the value of dynamic network information in identifying leukemia-related genes. The implementation of DyNDG is available at https://github.com/CSUBioGroup/DyNDG.

## Introduction

Blood disorders often give rise to pathological conditions that extend beyond the blood and may lead to dysfunction in other vital organs, posing a threat to human health. People may suffer from various types of blood conditions and blood cancers. Common blood disorders include anemia (such as hemolytic anemia), bleeding disorders (such as hemophilia, and blood clots), and blood cancers (such as leukemia, lymphoma, and myeloma). Leukemia, specifically, is a malignant blood disease characterized by the overproduction of immature or abnormal white blood cells, which eventually suppresses the production of normal blood cells and causes symptoms associated with cytopenias. According to the global cancer burden data released by the World Health Organization’s International Agency for Research on Cancer (IARC) in 2020, there were 9.96 million cancer deaths worldwide, including over 300,000 deaths from leukemia, while the number of leukemia deaths in China reached 60,000[1]. The occurrence and development of leukemia are closely linked to the mutation and abnormal expression of genes that regulate cell growth in the bone marrow [2], although genetic, behavioral, and environmental factors collectively contribute to leukemia [3]. Therefore, investigating the causative genes of leukemia holds significant importance to uncover its pathogenesis. Identifying the causative genes of leukemia is an extremely challenging task due to the pleiotropy of genes, the genetic heterogeneity of diseases, and the limited number of study subjects[4─6].

With the continuous explosion of biological data, computational methods for predicting potential pathogenic genes have emerged, playing a significant role in disease research, prevention, and detection[7]. The functions of biomolecular components in cells are often interdependent rather than acting independently. Diseases typically arise from disruptions in a complex biomolecular network caused by genetic variants, pathogens, and epigenetic changes, rather than solely from abnormal expression of individual genes[8]. Therefore, numerous classical disease-gene prediction methods based on biomolecular networks have been proposed [9 ─16], among which, network propagation has gained popularity due to its remarkable performance [17─21].

However, most traditional network-based methods are based on static biomolecular network structures. In contrast, cellular systems are highly dynamic and respond dynamically to external changes in different times, spaces, and conditions[22]. Thus, the relationships between biomolecules are constantly changing in response to time and conditions, and the molecular mechanisms underlying disease onset and progression are closely related to this dynamic nature. Traditional static networks lose the dynamic information due to their highly averaged and idealized structures, hindering the further advancement of network-based methods [23,24]. Some researchers have already shifted their attention from static biological networks to dynamic biological networks [25]. The key challenge in constructing dynamic biological networks is how to determine the dynamic features of biomolecule expression at different time points. Initially, De Lichtenbreg et al.[26] proposed that persistently expressed proteins are not dynamic, while cyclically expressed proteins are only expressed at their highest levels in the expression cycle. Hegde et al.[27] suggested using the average expression value of each region in the network as the threshold for judging which proteins are actually expressed in that region. Tang et al.[28] discovered that periodically expressed genes often exhibit peak expression levels greater than a fixed constant by studying the dynamical protein interaction network of yeast cells. In contrast to the former fixed threshold method, Zhang et al.[29] introduced a k-sigma method based on the 3-sigma rule by designing an activity threshold for each gene based on its dynamic expression. The development of dynamic protein network construction methods, which simulate the operational rules of real biological systems, effectively overcomes the limitations of static protein network-based analysis methods and plays a significant role in protein complex identification[30,31], protein function prediction[32,33], and biomarker identification[34,35]. Numerous studies have demonstrated that dynamic protein networks can yield superior results in various related issues since they can better reflect the dynamic properties of biological processes over time and in response to external environments[36,37]. It is necessary to simulate the dynamic changes of biomolecular networks during disease progression to reveal the associations between diseases and genes.

Therefore, we proposed the DyNDG model (Dynamic Network-based model integrating Differentially Expressed Genes) for identifying leukemia-related genes. We took three common leukemias: chronic myeloid leukemia (CML), chronic lymphocytic leukemia (CLL), and acute myeloid leukemia (AML) as research subjects. For each leukemia, DyNDG first generates a time-series dynamic network using expression data over stages of disease development, and then constructs a background-temporal multilayer network by integrating both the static network structure information and the dynamic information of leukemia development. Moreover, initialized by differentially expressed genes, DyNDG extends a random walk process into the multilayer network to extract scores of leukemia-related genes. Experimental results on the three types of leukemia and three control sets demonstrate that considering the time-series dynamics of leukemia development through a time-series dynamic network and a multilayer network framework significantly enhances the accuracy of prioritizing leukemia-related genes compared to popular approaches. Directly using predicted leukemia-related genes as drug targets may lead to interference and toxic side effects on normal cells. In order to minimize and avoid this possible harm, we selected higher-ranked genes from the predicted candidate gene list, exclude housekeeping genes from them and yield a set of promising candidate genes associated with leukemia. Through the integration of multiple analysis methods, we aimed to further understand the functions and regulatory mechanisms of these genes in the development of leukemia, providing valuable references and assistance for researchers and doctors.

## Method

### Datasets

#### Time-series expression data for leukemia

Gene Expression Omnibus database (GEO) is a public functional genomics data repository, which includes different gene expression datasets under different designs and conditions. Time series expression data for CML, CLL, and AML were obtained from the GEO database (https://www.ncbi.nlm.nih.gov/geo/; GEO accessions: GSE47927, [GSE2403](https://www.ncbi.nlm.nih.gov/geo/query/acc.cgi?acc=GSE47927), GSE122917), respectively. We preprocessed the data in the following three steps: 1) mapping of gene IDs and gene symbols; 2) filtering genes encoding proteins; 3) cleaning the data. Subsequently, the expression data were then subjected to pathological analysis to classify the stages of leukemia development.

The development of leukemia is a multi-factor and multi-step cancerous process. Leukemia is generally categorized into two types: acute leukemia and chronic leukemia, based on the differentiation and maturation status of leukemic cells and the natural progression of the disease [38]. AML, as a typical representative of acute leukemia, progresses rapidly and its disease stages are temporarily described as untreated, in remission, refractory, or recurrent[39] because there is currently no standard disease staging system for it. Chronic leukemia develops slowly and has a natural course of several years. In particular, CLL follows the Rai system, which classifies CLL into “low-risk group (stage 0), intermediate-risk group (stages I and II), and high-risk group (stages III and IV)” [40]. CML is categorized into three disease phases: chronic phase (CP), accelerated phase (AP), and blastic phase (BP) [41]. **Table 1** provides detailed information on the time series staging samples statistics for GSE47927, GSE2403, and GSE122917, based on their corresponding staging systems.

**Table 1.** The details of time series staging samples statistics.

| Disease | Data | Periods | Symbol | Samples |
| --- | --- | --- | --- | --- |
| CML | GSE47927 | Normal | normal | 15 |
|  |  | Chronic Phase | CP | 24 |
|  |  | Accelerated Phase | AP | 18 |
|  |  | Blastic Phase | BP | 10 |
| CLL | GSE2403 | Low-risk | Rai 0 | 8 |
|  |  | Intermediate-risk | Rai 1, Rai 2 | 7 |
|  |  | High-risk | Rai 3, Rai 4 | 6 |
| AML | GSE122917 | Normal | 0 | 3 |
|  |  | Untreated | 1 | 3 |
|  |  | Recurrent | 2 | 3 |

#### Biological networks

STRING is a database that covers the largest number of species and contains the most extensive information on protein-protein interactions. It collects and integrates known and predicted protein-protein association data from a variety of sources, including automated text mining of scientific literature, computational interaction predictions based on co-expression analysis, interaction experiments, and known complexes/pathways from curated sources[42]. HumanNet is an integrated human gene network database for disease research. It was constructed by incorporating new data types, expanding data sources, and utilizing improved network inference algorithms[43]. Due to the rich protein-protein interactions collected in STRING (https://cn.string-db.org/) and the focus of HumanNet (https://staging2.inetbio.org/humannetv3/) on disease research, we chose to use data from these two databases to construct the static protein-protein interaction (PPI) network separately. Further details regarding the static PPI networks can be found in Supplementary Table S1.

#### Disease genes for leukemia

Known leukemia-related genes were obtained from the MalaCards Database[44] (https://www.malacards.org/; MCID: [LKM071](https://www.malacards.org/card/leukemia_chronic_lymphocytic?search=CLL), [LKM063](https://www.malacards.org/card/leukemia_chronic_myeloid?search=CML), [LKM061](https://www.malacards.org/card/leukemia_acute_myeloid?search=AML)) which is an integrated database of human diseases and their annotations. There are 527 AML-related genes, 204 CML-related genes, and 339 CLL-related genes collected from the MalaCards database. We filtered these genes using protein-coding genes curated from the HUGO Gene Nomenclature Committee (HGNC) database[45] (https://housekeeping.unicamp.br/?download).

### Framework of DyNDG

Here, we proposed a dynamic network-based model, DyNDG, for predicting leukemia-related genes. DyNDG involves constructing the time-series dynamic biological networks and the background-temporal multilayer biological network, followed by network propagation on the multilayer biological network (see **Figure 1** for its schematic diagram). The original inputs consist of a gene expression matrix and a static PPI network (**Figure 1A)**. After constructing the time-series dynamic network (**Figure 1B**), the S-layer time-series dynamic network and the background network are combined to form the background-temporal multilayer biological network (**Figure 1C**). The initial probability vector of genes in the multilayer network is derived through differential gene expression analysis (see **Figure 1D** and the section “Network propagation on background-temporal multilayer biological network” for details). Subsequently, a network propagation process on the background-temporal multilayer biological network, which is initialized by differentially expressed genes, is performed to generate the predictive scores for leukemia-related genes (**Figure 1E**).

**Figure 1.**
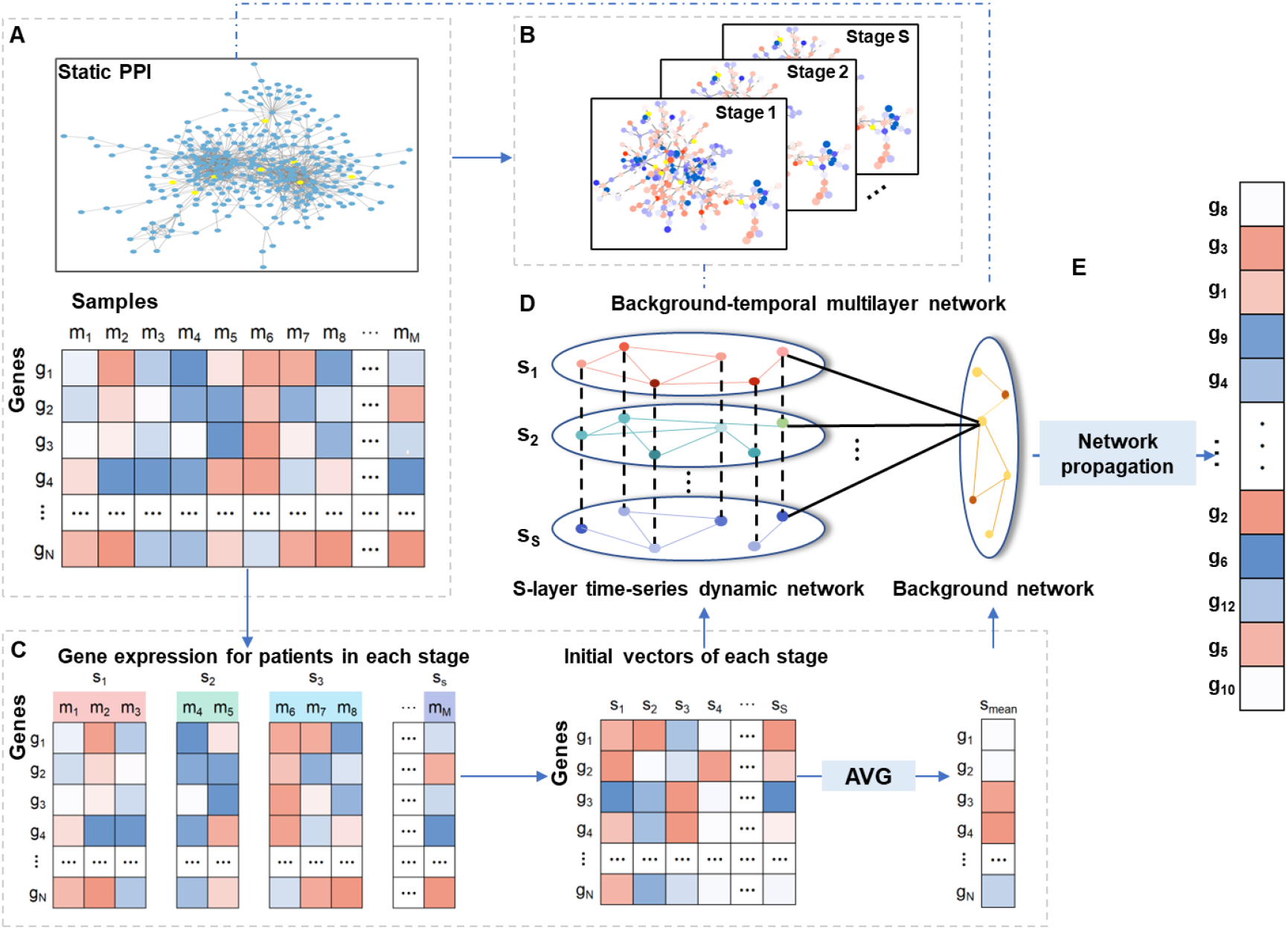
The schematic diagram of DyNDG A. The original inputs to DyNDG. **B**. The S-layer time-series dynamic network. **C**. Obtaining the initial probability vector of genes nodes for each stage through differential expressed genes analysis from time-series gene expression data of patients. The meaning of S_mean_ means taking the arithmetic mean of the initial probability vectors across all stages. S_mean_ is used as the initial probability vector of gene nodes in background network layer. **D**. The background-temporal multilayer biological network. The initial probability vector for each stage is applied to corresponding gene nodes in the corresponding-stage network layer. S_mean_ is applied to corresponding gene nodes in the background network layer. **E**. The vector of scores for leukemia-related genes is obtained as the output of network propagation on the background-temporal multilayer biological network.

### Construction of the time-series dynamic networks

The disease progression of leukemia involves multiple stages, and the biological networks associated with leukemia undergo dynamic changes throughout these stages. The gene expression data capturing the progression of leukemia was obtained by investigating patient populations at different stages of the disease, which can provide valuable insights into the dynamic gene changes that occur as leukemia progresses. The biological network constructed for each stage encompasses the gene interaction information from all patients within that stage. Given the inherent noise in gene expression data and the diverse expression patterns across genes, we employed the k-sigma method to calculate the active probability of each gene across various patient samples[29]. This approach effectively distinguishes the active level of genes in each patient sample when constructing the biological network for each disease stage. The gene expression matrix *G*^*N*×*M*^ contains expression profiles of *M* patient samples from *S* disease stages, where *N* is the number of genes. For each patient sample *m, G*_*m*_(*g*_*n*_) denotes the gene expression value of gene *g*_*n*_ in this patient. For all *M* patient samples, define 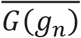 and *σ*(*g*_*n*_) as the algorithmic mean and standard deviation of gene expression values for *g*_*n*_ respectively. We calculated *k* -sigma threshold (where *k* represents the number of sigma) of each gene *g*_*n*_ by:

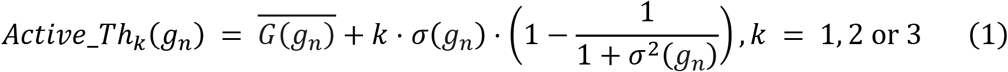

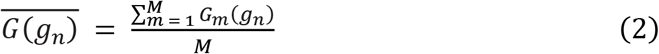

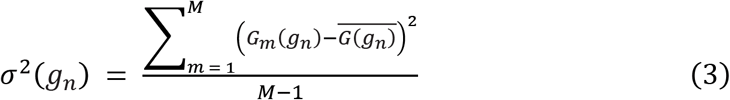

*Active_Th*_*k*_(*gn*) represents *k* active threshold of *g*_*n*_. The active probability of a protein corresponding to gene *g*_*n*_ in the patient sample *m* is calculated as follows [29]:

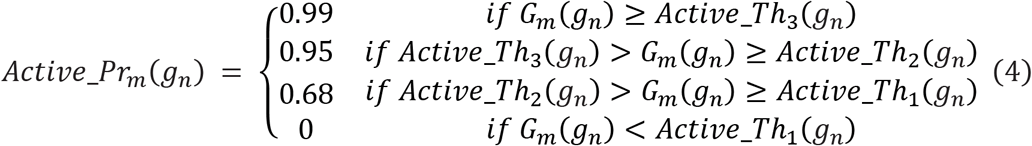

In general, the active probability value of a protein can be used as a measure of its active level in a patient sample. In order to construct the time-series dynamic network, we initially constructed the PPI network 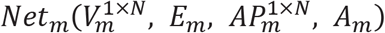 for each patient sample 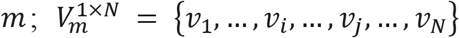 is the set of *N* gene/protein nodes; *E*_*m*_ is the set of interactions; 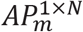 is the active probability vector of *N* gene/protein nodes; *A*_*m*_ represents the adjacency matrix which represents the confidence of the interactions between genes. Reweight *A*_*m*_ to 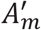 using 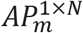 as follows:

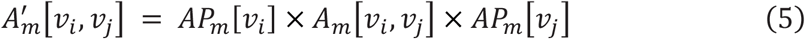

For the sake of concise and compact expression, a calculation method called “Broadcasting Multiplication” can be employed to describe the reweighting operation. The calculation process of Formula (5) can be found in section 1 of File S1.

Divide *M* samples into *S* leukemia stages, and the number of samples in each stage *s* is *M*_*s*_. The adjacency matrix *A*^[*s*]^(*s* = 1, … *S*) of leukemia stage *s* can be calculated as follows:

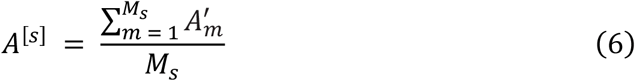

Finally, we constructed the time-series dynamic networks based on the adjacency matrix *A*^[*s*]^ of each leukemia stage. **Figure 2** shows the process of building the time-series dynamic networks. Firstly, the *k*-sigma threshold for each gene *g*_*n*_ is calculated from the gene expression matrix. The active probability vector of proteins corresponding to genes is obtained by using this threshold. By integrating the gene expression matrix and the static PPI network, we obtained the initial network for each patient sample. Further, new networks of patient samples are obtained by reweighting the edges with the active probabilities of the two corresponding genes using “Broadcasting Multiplication” (Broadcast Mul). The time-series dynamic networks corresponding to multiple leukemia stages is obtained by taking the “Union” of the patient sample networks. Note that the networks for all stages have the same number of genes, but the genes in different networks may be interconnected with different weights.

**Figure 2.**
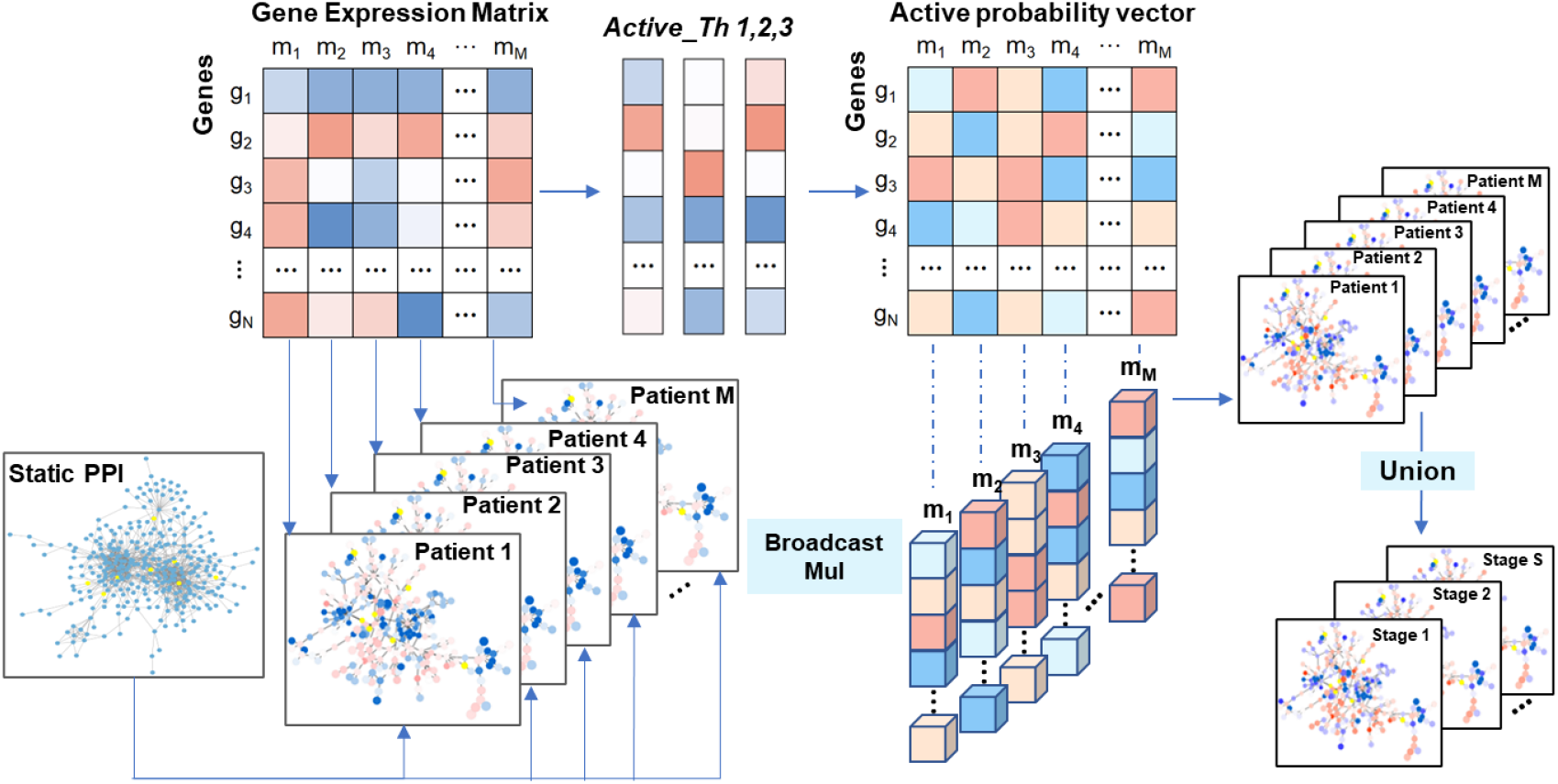
The schematic diagram of Time-series Dynamic Network construction. The process of constructing Time-series uynamic Network. Inputs consist of a gene expression matrix and a static PPI network. Union, network union of patient samples at the same stage.

### Construction of background-temporal multilayer biological network

The comprehensiveness and richness of biological networks significantly impact the ability to predict leukemia genes. To capture more realistic network changes throughout the course of leukemia, building upon the time-series dynamic networks simulating the dynamic alterations in biological networks, we proposed a background-temporal multilayer biological network framework. This framework integrates the dynamic networks and the static network into a multilayer network, effectively amalgamating information from both static and dynamic networks. The multilayer network is constructed by connecting shared gene nodes across different network layers, for jumping to a different layer through the same node when conducting propagation on it. The specific steps are as follows: Firstly, we constructed a *S*-layer temporal dynamic network by connecting the *S* time-series dynamic network layers of neighboring disease stages through shared nodes. Secondly, we connected the background network layer constructed by the static PPI network with each layer of the *S*-layer dynamic network by linking the corresponding gene counterparts. In the end, we obtained a connected multilayer network *Net*_*multi*_ = (*V*_*multi*_, *E*_*multi*_, *A*_*multi*_) with *L*(= *S* + 1) layers, where *S* is the number of layers in the above dynamic network; *A*_*multi*_ = {*A*^[1]^, … *A*^[*S*]^, *A*^[*L*]^} is the set of adjacency matrixes for the *L* layers, and *A*^[*L*]^ is the adjacency matrix of static PPI network; 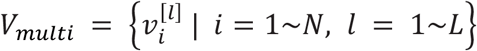 is the set of nodes (note that each gene has a counterpart node in each network layer); 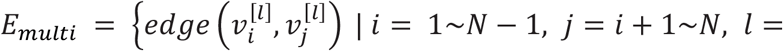
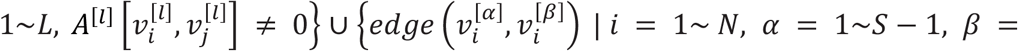
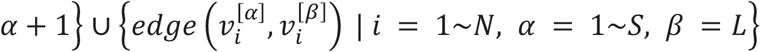 is the set of interlayer connections and intralayer PPIs. Note that all the edges here are undirected. Complex information about leukemia progression flows through edges in the multilayer network. The background-temporal multilayer biological network consists of two parts: the static network and the time-series dynamic network. The relevant details of the background-temporal multilayer biological network are presented in Supplementary Table S2.

### Network propagation on background-temporal multilayer biological network

To identify leukemia-related genes, we applied a network propagation process on the background-temporal multilayer biological network *Net*_*multi*_ During the walking process, the particles in the *Net*_*multi*_ either restart with a certain probability, or walk along a randomly selected (intra- or inter-layer) edge to a neighboring node (corresponding one of two possible actions: walk within the same layer, or jump to the counterparts of the same node in a different layer). Finally, as the random walk process concludes after a certain period, the scoring value of each node in the network will be obtained. During the random walk process, the scoring vector *P*_*t*+1_ of the nodes at step*t* + 1 can be obtained as follows:

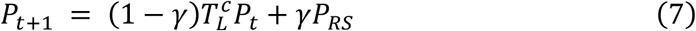

where *γ* ∈ (0,1) is the probability of restart, 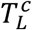 is the column-normalized transition matrix of the multilayer network, *P*_*RS*_ ∈ *R*^*NL*×1^ represents the initial scoring vector of *NL* nodes in the *L* layers network, *P*_*t*_ ∈ *R*^*NL*×1^ represents the scoring vector of *NL* nodes at step *t*.

Different stages of disease development are often accompanied by changes in gene expression levels and dynamic alterations in the network structure of gene interactions. However, most existing disease gene prediction methods do not take these biological phenomena into account. To address this, we proposed incorporating differential information during leukemia development as prior knowledge in the network propagation model. Specifically, for the gene expression matrix divided into *S* disease stages, we conducted differential gene expression analysis between each stage and the others by using the limma R package to obtain the T-statistic values of the genes in the corresponding layer of each stage. The absolute value of these T-statistic values is used to determine the initial scoring vector of genes, as they contain valuable information that effectively distinguishes each stage from the others. For the temporal multilayer network, we use the absolute T-statistics of the genes calculated in each of the S stages as the initial probability for each stage, thus incorporating stage-specific differential gene expression information. For the background network, we calculated the average of the above gene scores obtained from the differential analysis across all stages as the initial scores for each gene. By incorporating the T-statistic values into the background-temporal multilayer biological network, we can enhance the network with more comprehensive stage-specific information. As a result, we got the initial scoring vector *P*_*RS*_ ∈ *R*^*NL*×1^ in the *L*-layer network, which incorporates differential information of gene expression during disease progression as prior knowledge into the network.

Then, we constructed the *NL* × *NL* transition matrix *T*_*L*_ of the *L*-layer network by,

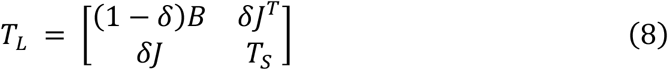

where *δ* ∈ [0, 1] *e* = (1,1, … …, 1)^*T*^ ∈ *R*^*S*×1^, 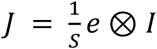 and *I* is a *N* × *N* identity matrix. Here, *B* is the adjacency matrix of the background network, *δ* controls the jumping probability of jumping between two types of network layers. *J* is the inter-network transition matrix that controls the jumping from the background network layer to each layer of the time-series dynamic network. Let *T*_*S*_ be the *NS* × *NS* transition matrix of the *S*-layer temporal dynamic network by,

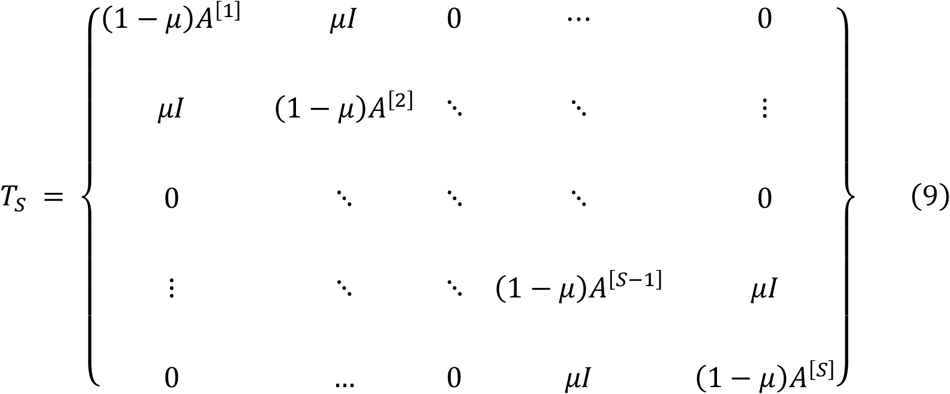

where *μ* ∈ [0, 1] controls the transition probability between the temporal dynamic network layers and *A*^[*s*]^ (*s* = 1 … … *S*) represents the adjacency matrix of each layer of the *S* -layer dynamic network. The matrix *T*_*S*_ controls the inter- and intra-layer jumps of the *S* -layer time-series dynamic network. We obtained 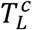 by column normalization of matrix *T*_*L*_. The example and parameter description of Formula (9) can be found in section 2 of File S1.

We plunged the column-normalized matrix 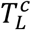 and initial scoring vector *P*_*RS*_ into the above iterative equation (7), which is iteratively updated until the difference between *P*_*t*_ and *P*_*t*+1_ falls below 10^−6^. Upon convergence, gene nodes in the background-temporal multilayer network are assigned a steady-state scoring vector *P*_∞_. In the background network, the steady-state scoring vector 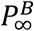 gives a measure of how strongly a gene is associated with leukemia. If 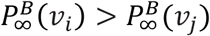, the gene corresponding to node *v*_*i*_ is more closely related to leukemia. Then genes are sorted in descending order based on 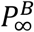, generating a ranking list of leukemia-related genes. Therefore, the higher-ranked genes can be considered as promising genes related to leukemia.

### Evaluation methods

#### Evaluation strategy and metrics

We conducted a systematic evaluation of the proposed DyNDG model, which involves constructing a background-temporal multilayer network and predicting leukemia-related genes on this network. The set of candidate genes consists of both a test gene set and a control gene set. The test set is composed of known disease genes. We considered three different schemes for constructing the control set. 1) Artificial Linked Interval Control Set (ALI): In this scheme, resembling linkage analysis or association studies, we selected 99 control genes for each test gene from the genes located closest to the test gene on the same chromosome. 2) Randomized Control Set (RC): This scheme mimics the scenario in exome sequencing studies. We randomly selected 99 control genes for each test gene from the entire genome. 3)Whole Genomic Control Set (WG): In this scheme, all genes in the network, excluding the known disease genes, were considered as control genes. For each leukemia, let *Set*_*t*_ denote the test gene set, let *Set*_*c*_ denote the control gene set, while *Set* = {*Set*_*t*_, *Set*_*c*_} represents the set of candidate genes. Combining the ranking list of genes obtained by DyNDG with *Set* can obtain the top-k ranking list *R*_*k*_ of candidate genes. We can quantitatively calculate four common metrics, *ie*., Topk_Precision, Topk_Recall, AUROC (Area Under the Receiver Operating Characteristic Curve), and AUPRC (Area Under the Precision-Recall Curve), to evaluate the proposed method.

#### Comparison algorithms

Network-based methods have gained significant popularity in predicting disease genes and have demonstrated remarkable prediction results. To compare the performance, we considered two categories of network-based disease gene prediction algorithms (see Supplementary Table S3 for details). For each category, we selected representative algorithms: 1) algorithms based on the single network: RWR[19]; 2) algorithms based on multiple networks: methods utilizing multiple independent networks such as RWRDRS[46], and the methods based on multiple interconnected networks like RWRMG[18] and RWRMP[20]. Furthermore, it is a common strategy to leverage the differences in gene expression levels between normal and disease stages in predicting disease genes. Hence, we also included two representative algorithms: 1) the T-test method, which generates disease association measures similar to network-based methods but without utilizing network information. 2) the DISNEP method, which enhances a general gene network into a disease-specific network and then prioritizes gene associations on the enhanced network[47].

### Functional enrichment analysis

Functional enrichment analysis is a technique widely used in bioinformatics to identify biological functions or processes that are over-represented in a given set of genes or proteins. This analysis helps in understanding the biological significance of a gene list or a set of differentially expressed genes by linking them to known biological functions, pathways, or processes. We used the clusterProfiler R package to perform GO enrichment analysis and KEGG pathway enrichment analysis on the top 1% ranked genes in the predicted leukemia-related candidate gene list to investigate their associations with leukemia-related functions and metabolic pathways.

## Results and discussion

### DyNDG can greatly improve predictive ability

To evaluate the performance of DyNDG quantitatively, we generated the set of leukemia-related candidate genes by the decreasing order of the predictive scores, and the evaluation of DyNDG was conducted by four common metrics (ie., Topk_Precision, Topk_Recall, AUROC, and AUPRC), along with three kinds of the control sets: ALI, RC and WG.

In identifying leukemia-related genes, our DyNDG method significantly outperforms the single-network-based method RWR[19] based on the static network upon Topk_Recall and Topk_Precision metrics, because of incorporating temporal dynamic network information of disease progression, while RWR demonstrates better performance than the classical T-test method (see **Figure 3** for details). Multi-network-based methods have been designed to operate on multiple independent or interconnected networks, enabling effective utilization of more information from these networks. Here, we also compared our DyNDG method with three Multi-network-based methods: RWRDRS[46], RWRMG[18], and RWRMP[20]. The comparison results in **Figure 4A-C** respectively show that DyNDG outperforms these Multi-network-based methods in identifying CLL-related genes, CML-related genes, and AML-related genes, in terms of Topk_Recall performance. It can be observed that increasing the k value appropriately leads to a noticeable improvement in Topk_Recall. It is indeed a difficult task to search for disease-related genes in almost full gene set. The recall scores of all other algorithms are also relatively low. Additionally, if the candidate range for search/prediction is limited, the recall of the algorithm will usually improve, including utilizing more known information. The results in **Figure 4D-F** also demonstrate that DyNDG can more precisely predict CLL-, CML-, and AML-related genes, respectively, in terms of Topk_Precision performance. Supplementary Figure S1 shows the overlaps of the top 100 genes predicted by different comparison methods for CLL, CML, and AML. Upon analyzing the overlaps of candidate genes predicted by DyNDG and comparison methods, we found that there is more than 50% overlap between DyNDG and the comparison methods, with the majority of the overlapping predictions coming from Multi-network methods. Additionally, DyNDG recommended some genes that were not identified by other algorithms which further illustrates the potential of DyNDG to complement and enhance existing predictive approaches. Supplementary Figure S2 ─S5 also show that DyNDG achieved the best on three leukemia datasets and different kinds of the control sets upon Topk_Recall and Topk_Precision, re-affirming that DyNDG can obtain more reliable and stable results, independent to the types of control sets.

**Figure 3.**
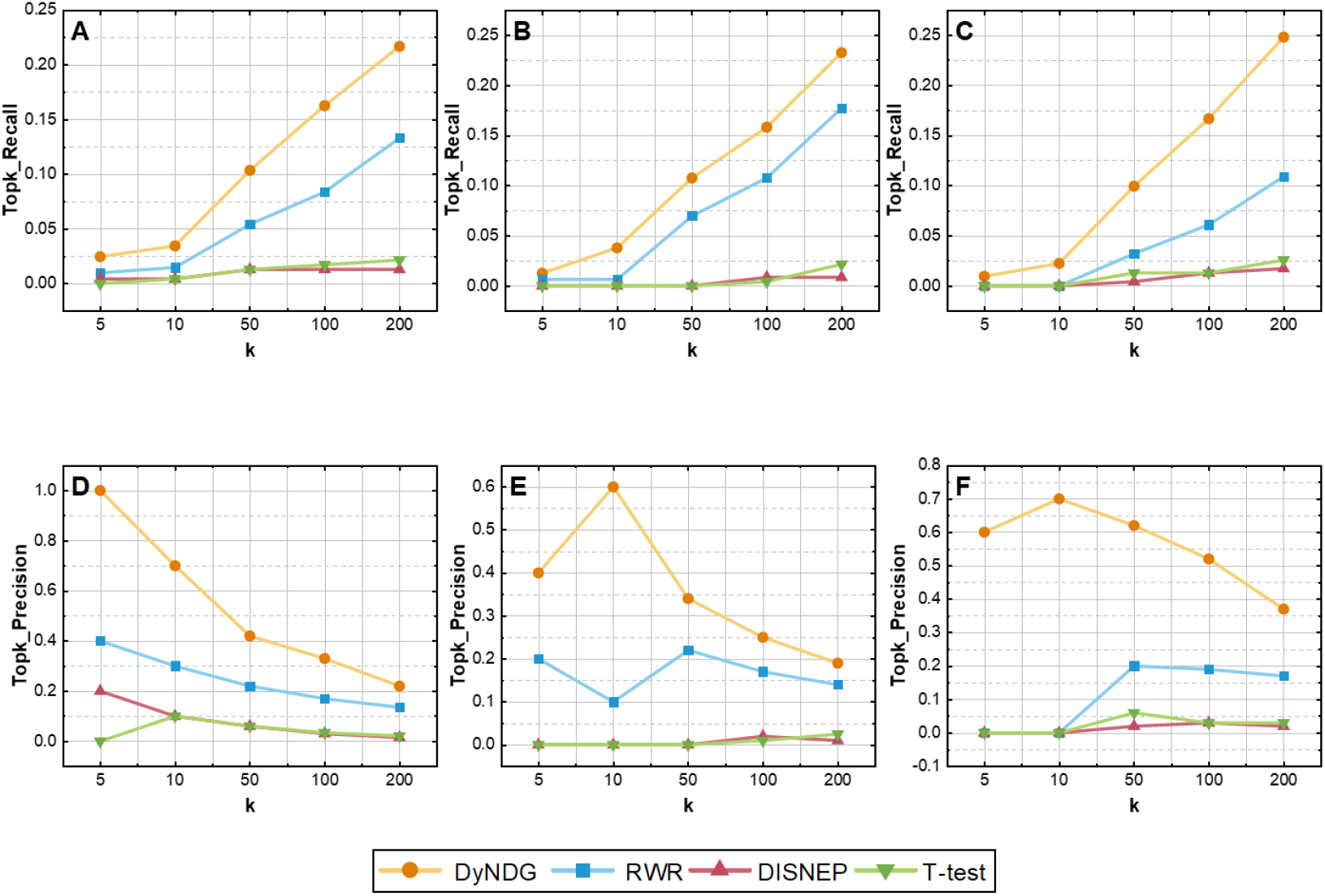
Performance comparison of DyNDG, RWR, T-test, and DISNEP under the WG control set, in term of Topk_Recall and Topk_Precision. **A, B**, and **C** present the Topk_Recall results for predicting CLL-, CML-, and AML-related genes, respectively. Similarly, **D, E**, and **F** display the Topk_Precision results for predicting CLL-, CML-, and AML-related genes, respectively. When k is 5, 10, 50, 100, and 200, DyNDG consistently exhibits higher Topk_Recall and Topk_Precision values than other methods.

**Figure 4.**
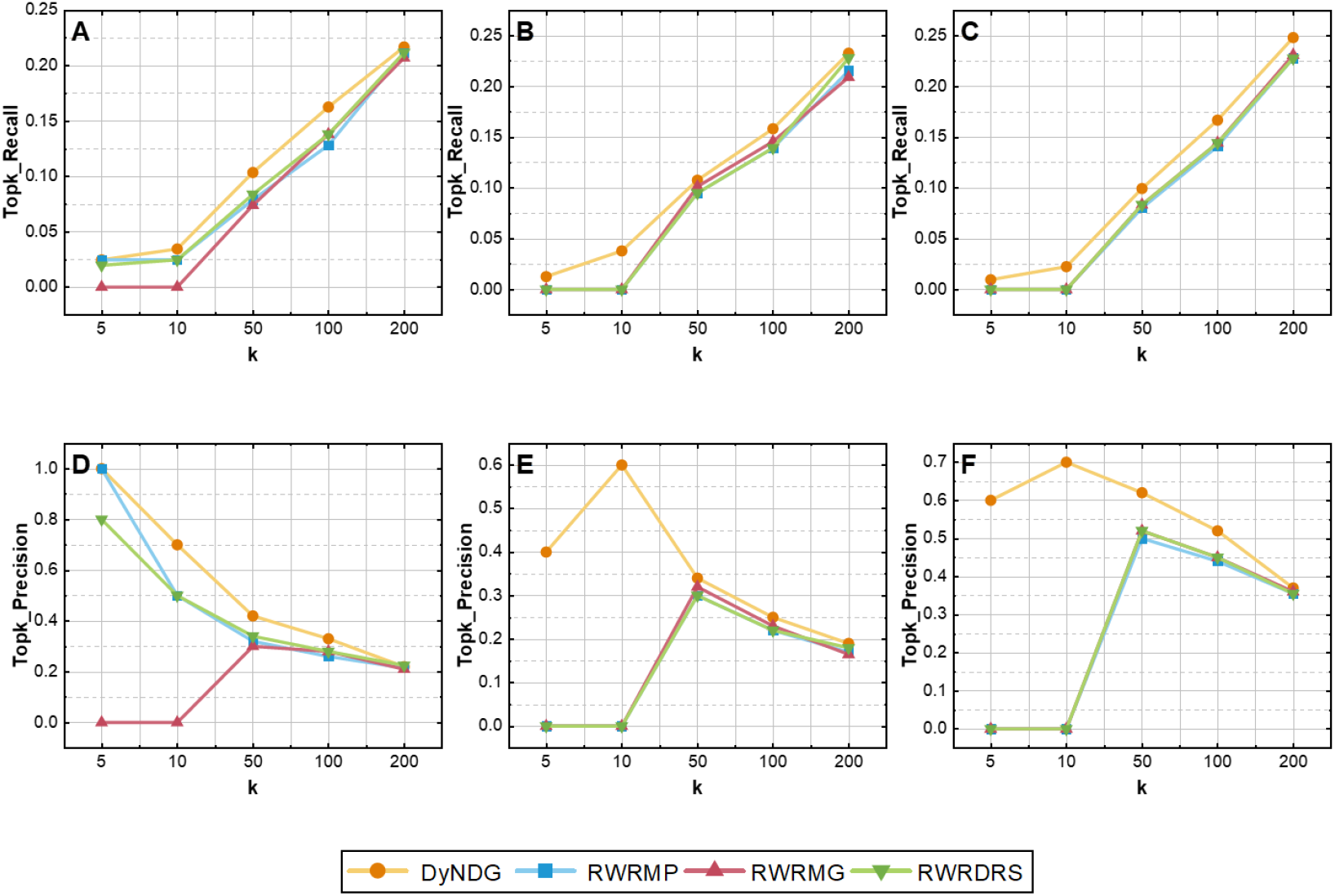
Performance comparison of DyNDG, RWRDRS, RWRMG, and RWRMP under the WG control set, in term of Topk_Recall and Topk_Precision. **A, B**, and **C** illustrate the Topk_Recall results for predicting CLL-, CML-, and AML-related genes, respectively. **D, E**, and **F** display the Topk_Precision results for predicting CLL-, CML-, and AML-related genes, respectively. When k is 5, 10, 50, 100, and 200, DyNDG consistently outperforms other methods in terms of Topk_Recall and Topk_Precision.

Moreover, the results of the performance comparison between DyNDG and other methods, using the AUROC and AUPRC metrics, are shown in **Table 2**. DyNDG significantly outperforms all the comparison algorithms in terms of AUROC and AUPRC, demonstrating a notable advancement in identification of leukemia disease genes.

**Table 2.** Performance comparison of DyNDG and other methods under AUROC and AUPRC metrics.

| Leukemia | Type | Methods | AUROC | AUPRC |
| --- | --- | --- | --- | --- |
| CLL | Single-network | RWR | 0.8080 | 0.0646 |
|  |  | RWRMP | 0.7967 | 0.1396 |
|  | Multi-network | RWRMG | 0.8394 | 0.1078 |
|  |  | RWRDRS | 0.7941 | 0.1433 |
|  | Differential gene | T-test | 0.7116 | 0.0224 |
|  |  | DISNEP | 0.7177 | 0.0231 |
|  |  | **DyNDG** | **0.8875** | **0.1753** |
| CML | Single-network | RWR | 0.7881 | 0.0676 |
|  | Multi-network | RWRMP | 0.8487 | 0.0910 |
|  |  | RWRMG | 0.8308 | 0.0920 |
|  |  | RWRDRS | 0.8556 | 0.0940 |
|  | Differential gene | T-test | 0.5798 | 0.0162 |
|  |  | DISNEP | 0.5712 | 0.0154 |
|  |  | **DyNDG** | **0.8983** | **0.1260** |
| AML | Single-network | RWR | 0.7493 | 0.0745 |
|  | Multi-network | RWRMP | 0.8422 | 0.1887 |
|  |  | RWRMG | 0.8445 | 0.1918 |
|  |  | RWRDRS | 0.8433 | 0.1927 |
|  | Differential gene | T-test | 0.5478 | 0.0155 |
|  |  | DISNEP | 0.5402 | 0.0145 |
|  |  | **DyNDG** | **0.8797** | **0.2334** |

### Effect of parameters and data

We systematically examined the impact of parameters (*ie*., *δ μ* and *γ*) on the performance of the DyNDG model in three types of leukemia. Details of the analysis can be found in section 3 of File S1, and the results are presented in Supplementary Figure S6 ─ S11. *δ* and *μ* regulate the transitions of the background-temporal multilayer network. The results indicate that the model performance remains relatively stable as the parameters *δ* and *μ* vary, which suggests that the multilayer network framework can effectively integrate background network knowledge and temporal dynamic network information, ensuring the robustness of the model. A lower restart probability *γ* corresponds to better model performance, indicating that the background-temporal multilayer network contains rich information related to leukemia disease genes and is crucial for predictive performance. Therefore, we chose *δ* = 0.5, *μ* = 0.5, *γ* = 0.1 as the parameter combination that yields the relatively best model performance.

Additionally, we investigated the influence of different static network data on the performance of DyNDG. Further information about this study can be found in section 4 of File S1. The results in Supplementary Figure S12 show that DyNDG performs better when using the static PPI network of STRING[42] which provides a broader and more diverse range of PPI information. Based on Supplementary Figure S13 and S14, we also studied the differences in the initial probability distribution between different layers of the background-temporal multilayer network. The initial probabilities of the background network comprehensively consider the characteristics of the temporal multilayer networks at various stages. The differences in the initial probability distribution across different stages of disease progression indicate that the T-statistic distributions obtained from differential gene analysis at each stage can effectively represent the characteristics of each stage.

To explore how different components of the DyNDG model contribute to enhancing the predictive power, we evaluated two different experiments: (1) DyNDG_static, where the random-walk process was performed only in the background network; (2) DyNDG_dynet, where the random-walk process was conducted exclusively in the dynamic network (The detailed experimental setup can be found in section 5 of File S1). The experimental results (Supplementary Figure S15) show that the background network plays a crucial role in the model which builds upon universal interaction patterns in biomolecules, providing a comprehensive background and a solid foundation for our model. Simultaneously, the introduction of the dynamic network enhances the predictive capability of the model, particularly in simulating dynamic changes in biological networks at different leukemia stages and capturing potential gene associations. This suggests that in the prediction of leukemia-related genes, the collaboration of static and dynamic networks contributes to the overall improvement in predictive performance of model.

### Comprehensive analysis of candidate disease genes for AML, CLL, and CML

In this study, we further conducted case studies and comprehensive analysis for specific leukemia (*ie*. AML, CLL, and CML). Specifically, we applied DyNDG to each of the three leukemias and obtained a ranked list of genes for each of them. Then, we individually filtered each of the three gene lists, by removing the genes reported in Malacards[44] that are associated with the respective types of leukemia. Additionally, we excluded the human housekeeping genes collected from the HRT Atlas database[48]. As a result, we obtained three lists of candidate genes associated with their respective types of leukemia.

#### Integrative analysis of function annotations and clinical survival in AML

AML is the most common and lethal adult acute leukemia, characterized by its aggressive nature, poor prognosis, and high susceptibility to relapse after treatment[49]. Through the application of DyNDG, we can identify AML-related genes that have the potential to drive AML progression and contribute to poor prognosis in patients.

Functional enrichment analysis conducted on the top 1% ranked genes in the AML-related candidate gene list revealed the KEGG pathways and GO terms most associated with AML, as shown in **Figure 5A** and **Figure 5B** respectively. The most relevant KEGG pathways consist of cancer-related pathways (e.g., cell cycle and MAPK signaling pathway), signaling pathways regulating pluripotency of stem cells, and leukemia-related pathways (e.g., leukocyte transendothelial migration and platelet activation). The GO enrichment analysis shows many AML-related biological processes (e.g., leukocyte cell-cell adhesion, leukocyte proliferation and cellular response to inorganic substance), molecular functions (e.g., catalytic activity, acting on DNA and cyclic nucleotide-dependent protein kinase activity) and cellular components (e.g., chromosomal region). To further investigate the candidate genes enriched in these AML-related pathways, we obtained a set of multi-pathway enrichment genes as shown in **Figure 5C** (The corresponding relationships between GO term IDs and Descriptions, as well as between KEGG pathway IDs and Descriptions, can be found in Supplementary Table S4).

**Figure 5.**
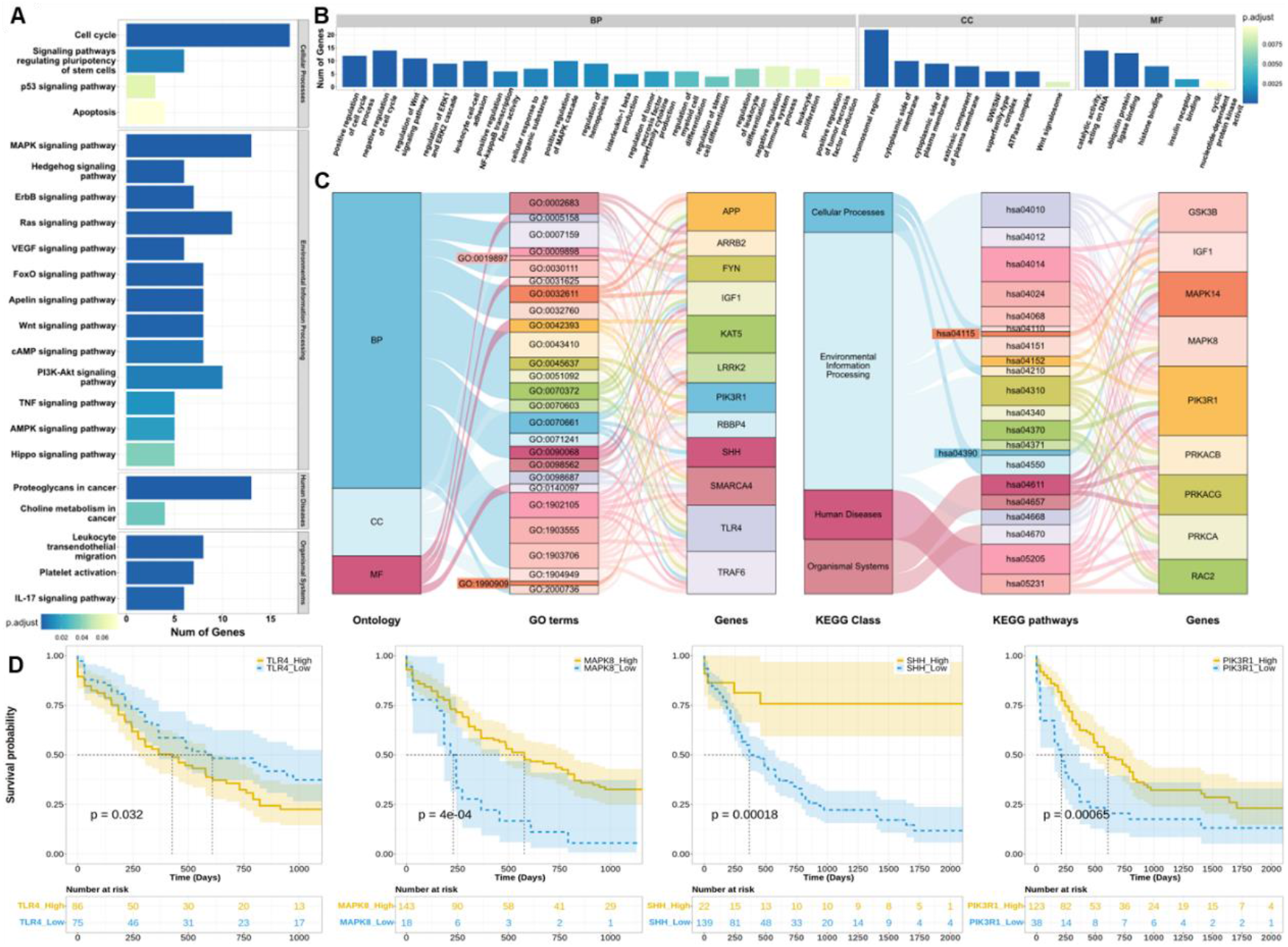
DyNDG predicts AML-related genes that are closely associated with clinical survival. **A**. The bar chart showcases the results of KEGG pathway enrichment analysis, highlighting the most relevant KEGG pathways associated with AML. **B**. The bar chart illustrates the outcomes of GO enrichment analysis, highlighting the most relevant GO terms associated with AML. **C**. The Sankey diagrams show the associations between GO terms and multi-pathway enriched genes, as well as the associations between KEGG pathways and multi-pathway enriched genes. **D**. Kaplan–Meier overall survival curves of AML patients grouped by the averaged expression of *TLR4, MAPK8, SHH*, and *PIK3R1*, respectively (with the median value as the threshold). The P value was calculated by the log-rank test.

By conducting survival analysis as shown in **Figure 5D**, we observed significant differences in overall survival among patient groups with different expression levels of many multi-pathway-enriched genes (e.g., *TLR4, MAPK8, SHH*, and *PIK3R1*). *TLR4* is expressed by acute myeloid leukemia cells, several bone marrow stromal cells, and non-leukemic cells involved in inflammation. *TLR4* in leukemia cells is of significant importance for the growth and development of leukemia cells in human AML, and targeting *TLR4* may have direct and indirect effects on leukemogenesis. A study has indicated that high expression of *TLR4* is associated with a decrease in survival rates after intensified anti-leukemia treatment[50]. This suggests that high expression of *TLR4* may be related to lower survival rates in AML patients, which is consistent with our survival analysis results of *TLR4* shown in **Figure 5D**. *MAPK8*, also known as *JNK1*, is a member of the MAPK family. The aberrant activation of the MAPK signaling pathway is closely associated with the proliferation, survival, drug resistance, and metastatic capacity of AML cells[51]. Therefore, as a member involved in regulating the MAPK signaling pathway, *MAPK8* may play an important role in key biological processes regulating AML. Studies have shown that in AML, certain anticancer drugs may exert their anticancer effects by activating *MAPK8* to inhibit cell proliferation in AML cell lines[52]. In the clinical survival analysis presented in **Figure 5D**, it was observed that low expression of *MAPK8* may be associated with lower survival rates. This could be attributed to the aberrant activation or impaired function of the MAPK signaling pathway resulting from the lower expression of *MAPK8*, ultimately leading to reduced survival rates in AML patients. These findings provide important clues for further investigating the role of *MAPK8* in the development and treatment of AML. *SHH* has been implicated in the maintenance of cancer stem cells (CSCs), playing a critical role in the development of drug resistance and disease relapse in AML. Research findings suggest that *SHH* could serve as a prognostic marker for CSCs in AML, providing valuable insights into disease outcomes[53]. Furthermore, in the presence of LPS/TNF-α/IFN, *SHH* antagonists have been utilized for the treatment of AML patients[54]. Previous evidence suggests that *SHH* may be associated with the occurrence, progression, and treatment of AML. As shown in **Figure 5D**, a significant difference in survival probability was observed between patient groups with different levels of *SHH* expression, providing clinical evidence for the pivotal role of *SHH* in the development of AML. *PIK3R1* is the gene encoding the p85α subunit of the PI3K regulatory subunit, playing a critical regulatory role in the PI3K/Akt signaling pathway. Previous studies have indicated that *PIK3R1* is an actionable gene in AML[55]. The PI3K/Akt signaling pathway regulated by *PIK3R1* is involved in various cellular processes such as cell survival, proliferation, and metabolism in normal cells. Its aberrant activation is associated with the development and progression of multiple tumor types, including AML. In 50─80% of AML patients, constitutive activation of the PI3K/Akt pathway has been detected, which is associated with reduced overall survival[56]. Therefore, the abnormal expression of *PIK3R1*, which regulates the PI3K/Akt pathway, may be implicated in the occurrence and progression of AML. **Figure 5D** illustrates the relationship between *PIK3R1* expression and the survival rate of AML. The identification of these genes provides valuable insights into the potential molecular mechanisms, pathogenesis, and key factors influencing the prognosis of AML. It offers opportunities to identify new therapeutic targets and has the potential to improve prognosis assessment and develop personalized treatment strategies.

#### Integrative analysis of function annotations and network in CLL

In Western countries, CLL is one of the most prevalent types of leukemia, and there is currently no definitive cure available. Despite advancements in treatment with novel therapies and targeted drugs, there is a continuing need for new targeted treatments and innovative strategies to improve efficacy and survival rates[57]. By applying DyNDG to CLL, we can predict CLL-related genes that can serve as references for discovering new therapeutic targets and developing personalized treatment strategies.

Functional enrichment analysis conducted on the top 1% candidate genes revealed the KEGG pathways and GO terms most relevant to CLL, as shown in **Figure 6A** and **Figure 6B** respectively. The KEGG pathways most relevant to CLL include cancer-related pathways (such as cell cycle, MAPK signaling pathway, and transcriptional misregulation in cancer), B cell receptor signaling pathway, and pathways associated with chronic leukemias (such as chronic myeloid leukemia and leukocyte transendothelial migration). GO enrichment analysis revealed numerous CLL-related biological processes (such as leukocyte cell-cell adhesion, leukocyte proliferation, lymphocyte proliferation, and regulation of tumor necrosis factor production), molecular functions (such as kinase regulatory activity, SMAD binding, and G protein-coupled receptor binding), and cellular components (such as chromosomal regions). Then, we obtained a set of multi-pathway enriched genes, which are enriched in multiple GO terms and KEGG pathways, through Sankey diagrams as shown in **Figure 6C** (Enriched pathways are identified by their respective GO term IDs and KEGG pathway IDs, which can be found in Supplementary Table S4, along with their descriptions).

**Figure 6.**
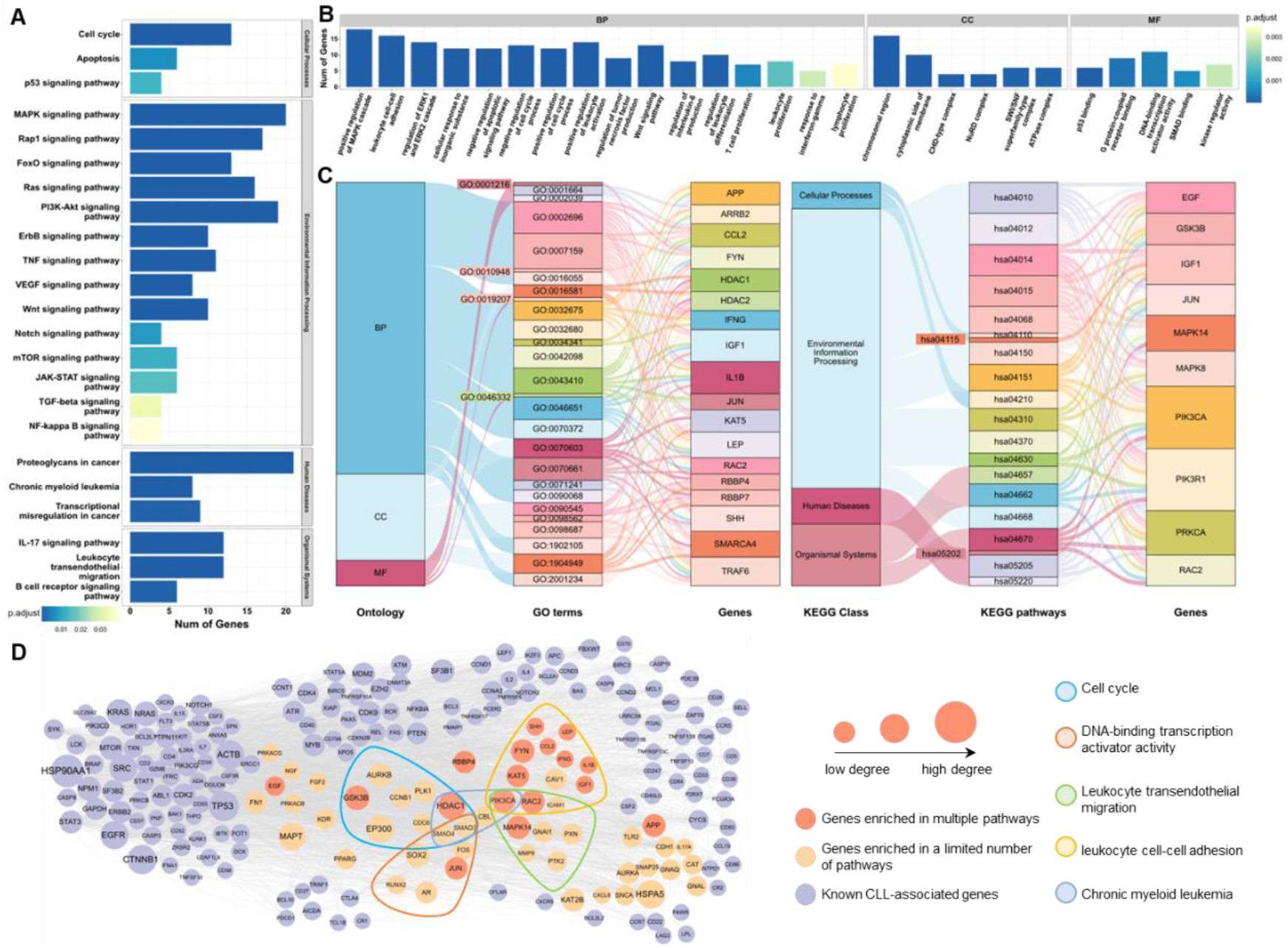
DyNDG predicts CLL-related genes that possess network hub characteristics. **A**. The bar chart showcases the results of KEGG pathway enrichment analysis, highlighting the most relevant KEGG pathways associated with CLL. **B**. The bar chart illustrates the outcomes of GO enrichment analysis, highlighting the most relevant GO terms associated with CLL. **C**. The Sankey diagrams show the associations between GO terms and multi-pathway enriched genes, as well as the associations between KEGG pathways and multi-pathway enriched genes. **D**. The network graph reflects the degree centrality of candidate genes enriched in CLL-related pathways and known CLL-related genes.

To explore the criticality of candidate genes enriched in CLL-related pathways in the PPI network, we conducted network analysis using Cytoscape[58]. Based on human static PPI network sourced from the STRING database[42], we analyzed the node degree centrality of candidate genes enriched in CLL-related pathways and CLL-related genes reported in Malacards[44]. Subsequently, we generated a network diagram as shown in **Figure 6D**. It is well-known that genes with high degree centrality play a crucial role in regulating and influencing information transmission, network stability, functional integration, and disease relevance within the PPI network. In the static PPI network, candidate genes enriched in CLL-related pathways exhibit degrees similar to known CLL-related genes. This suggests that the candidate genes may have similar network influence as the known CLL-related genes and could play a crucial role in the progression of CLL. Many of these candidate genes even surpass certain known CLL-related genes in terms of degree centrality, indicating that they may have a greater importance in regulating the network than some of the known key genes. *HDAC1, PIK3CA*, and *RAC2* are all genes that are enriched in multiple pathways and exhibit high degree centrality in the PPI network. Lastly, we conducted literature research on *HDAC1, PIK3CA*, and *RAC2. HDAC1* is one of the components of the histone deacetylase complex, and elevated *HDAC* enzyme activity has been found to be associated with the development of leukemia and other cancers. Significantly elevated levels of *HDAC1* expression have been observed in CLL[59], and studies have indicated that *HDAC1* can act as a transcriptional activator in CLL, promoting CLL cell survival and progression[60]. *PIK3CA* is a component of the PI3K pathway and has long been described as an oncogene. Inhibition of PI3K subtype associated with *PIK3CA* has been shown to be potentially valuable in CLL[61]. *RAC2* is a member of the *RAC* subfamily of Rho GTPases and is closely associated with oncogenic signaling. Existing studies have found that *RAC2* can activate two known PLC*γ*2 mutations in CLL, suggesting that *RAC2* may play a significant role in the survival and proliferation of CLL cells carrying these mutations[62]. Research has also indicated that upregulation of *RAC2* expression may be associated with resistance to ibrutinib in CLL[63]. In conclusion, *HDAC1, PIK3CA*, and *RAC2* may have significant biological functions and clinical implications in CLL. They could potentially serve as potential therapeutic targets or biomarkers for predicting CLL progression and treatment response.

#### Integrative analysis of function annotations and differential expression in CML

CML is a chronic leukemia caused by the fusion of the BCR-ABL genes. Inhibiting the activity of the BCR-ABL fusion protein with tyrosine kinase inhibitors (TKIs) can effectively control the proliferation and survival of CML cells, leading to sustained clinical and molecular remission in the majority of patients. However, long-term use of TKIs may result in side effects, and some patients may develop resistance or intolerance to TKI treatment[64]. By applying DyNDG to CML, predicting genes associated with its disease progression may provide valuable insights for studying new targets in CML, addressing the issue of drug resistance, and exploring novel therapeutic approaches for CML. Functional enrichment analysis conducted on the top 1% candidate genes revealed the KEGG pathways and GO terms most related to CML, as shown in **Figure 7A** and **Figure 7B** respectively.

**Figure 7.**
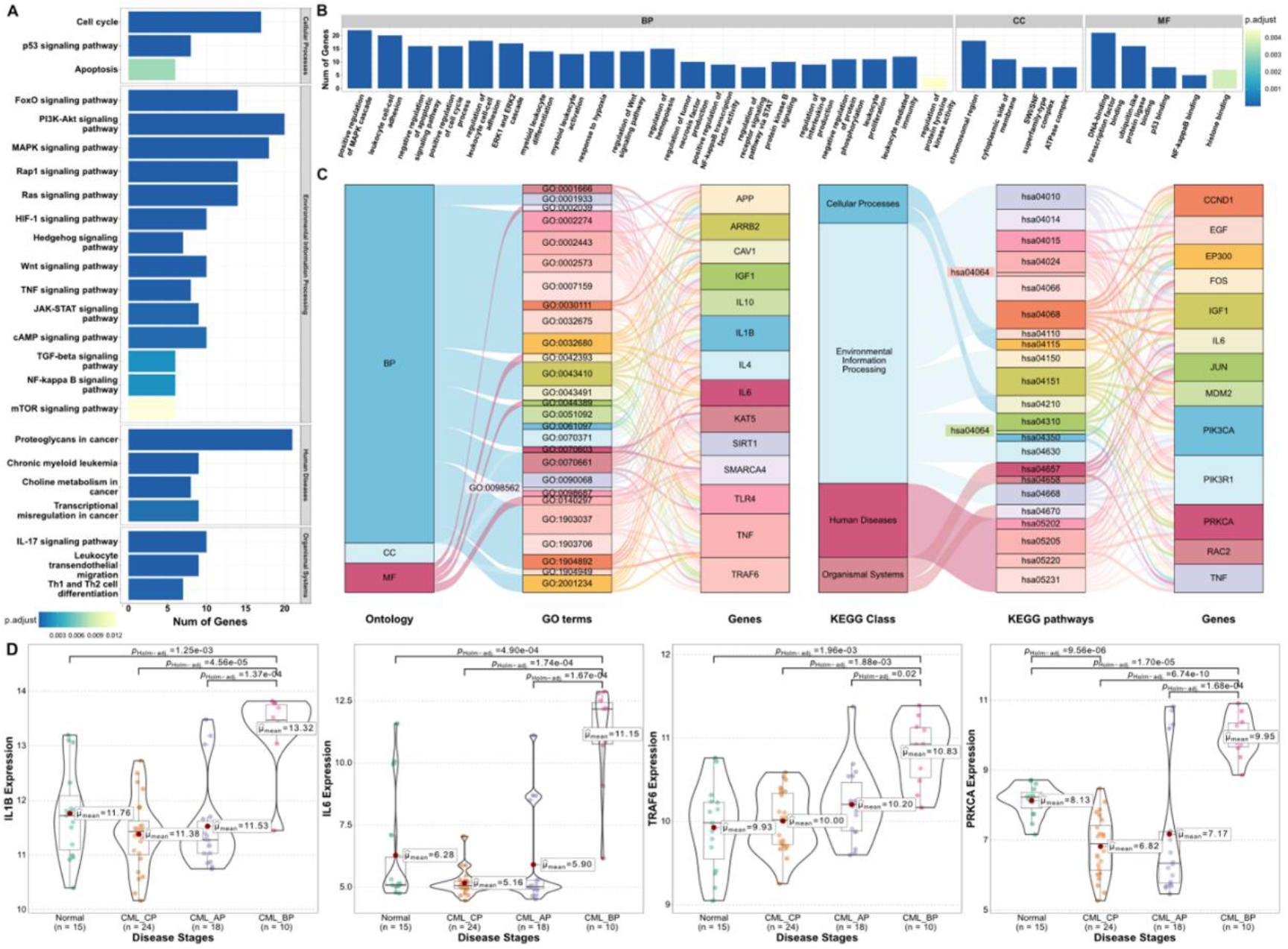
DyNDG predicts CML-related genes that are closely associated with disease progression. **A**. The bar chart showcases the results of KEGG pathway enrichment analysis, highlighting the most relevant KEGG pathways associated with CML. **B**. The bar chart illustrates the outcomes of GO enrichment analysis, highlighting the most relevant GO terms associated with CML. **C**. The Sankey diagrams show the associations between GO terms and multi-pathway enriched genes, as well as the associations between KEGG pathways and multi-pathway enriched genes. **D**. The violin plots display the significant differences between CML disease progression stages for *IL1B, IL6, TRAF6*, and *PRKCA*, respectively.

The KEGG pathways most relevant to CML include cancer-related pathways (such as cell cycle, PI3K-Akt signaling pathway, and proteoglycans in cancer), the pathway closely associated with the immunity (such as Th1 and Th2 cell differentiation), and pathways associated with CML (such as chronic myeloid leukemia and leukocyte transendothelial migration). GO enrichment analysis revealed numerous CML-related biological processes (such as leukocyte cell-cell adhesion, leukocyte proliferation, myeloid leukocyte differentiation, and leukocyte mediated immunity), molecular functions (such as DNA-binding transcription factor binding, and ubiquitin-like protein ligase binding), and cellular components (such as chromosomal regions). In order to identify key genes for further research, we generated Sankey diagrams, as shown in **Figure 7C**, to obtain genes enriched in multiple GO terms and KEGG pathways (The IDs and corresponding descriptions of GO terms and KEGG pathways can be found in Supplementary Table S4).

Then, studying the expression level changes of these key genes in different pathological stages of CML, we found significant differences in the expression of many genes across different stages of CML development. Specifically, compared to other stages, *IL1B, IL6, TRAF6*, and *PRKCA* show significantly higher expression levels in the CML_BP stage as shown in **Figure 7D**, indicating their association with CML disease progression and their potential as specific markers for the late-stage of CML. The extensive previous research can provide support for our analysis. *IL1B* is an important member of the IL-1 family and serves as a crucial cytokine involved in various inflammatory processes and certain anti-tumor physiological mechanisms. The increase in *IL1B* levels may promote CML resistance to tyrosine kinase inhibitors (TKIs) by enhancing cell viability and facilitating cell migration[65]. Recent studies have shown that the development of CML is accompanied by elevated levels of *IL1B* [66]. *IL6*, a pleiotropic cytokine, plays an important role in some processes associated with leukemia such as immune response and hematopoiesis. *IL6* has been regarded as a valuable prognostic marker in CML and may be correlated with the transformation of different phases in CML because *IL6* expression in CML-BP is significantly upregulated compared with in CML-CP and CML-AP[67]. *TRAF6* is a member of the superfamily of TRAF proteins. It is not only overexpressed in many cancer tissues but also closely associated with tumor cell proliferation, migration, and apoptosis. Studies have shown that interfering with or inhibiting the role of *TRAF6* in tumor-related signaling pathways may provide new therapeutic approaches for cancer treatment[68]. In patients with CML undergoing imatinib treatment, downregulation of *TRAF6* leads to higher levels of cellular apoptosis, indicating the significant regulatory role of *TRAF6* in CML cell survival and treatment response[69]. *PRKCA*, also known as *PKCα*, is an important subtype of the Protein Kinase C (PKC) family. Aberrant regulation of different PKC subtypes is associated with the development of many human diseases. In CML cells, *PRKCA*, as a classical PKC isoform, exhibits significantly reduced kinase activity. This indicates that *PRKCA* may be functionally impaired or inactivated in CML, and it could contribute to abnormal proliferation and pathological development of CML cells[70].

### Validation by RNAiCRISPR in different types of cell lines of AML, CLL, and CML

To verify the effectiveness of our method DyNDG, we utilized the independent DepMap database[71] and investigated the roles of predicted candidate genes in different types of leukemia cell lines. Specifically, we analyzed three types of leukemia cell lines: AML, CLL, and CML, focusing on the top ten genes in our predicted candidate gene list. Notably, the “Gene Effect” values for most of these genes are below 0, with some even lower than -1, as shown in **Figure 8A, B and C**, which suggests that the majority of the top ten predicted genes are likely essential in the corresponding leukemia cell lines, as knocking out or down these genes can affect leukemia cell proliferation. Almost all of the top ten predicted genes can find at least one type of leukemia cell line where the knocking out or down of these genes shows significant affects. Some of them have been investigated in the diagnosis and prognosis of leukemia. For example, it has been shown that high expression of *HSPA8* is often associated with poorer survival rates in AML patients[72]; *HDAC1* can activate driver genes in CLL, thereby promoting the survival and progression of CLL[60]; *EP300* is a promising tumors therapeutic target. The mutations of *EP300* frequently occur in various types of hematologic malignancies including leukemia such as primary and relapsed pediatric ALL, aggressive NK-cell leukemia (ANKL), and CML-CP through a variety of different mechanisms[73]. These findings not only confirm the efficacy of our predictive method but also highlight the potential of these predicted candidate genes as therapeutic targets. Moreover, we examined the “Gene Effect” of known disease genes for AML, CLL, and CML in the DepMap database, as shown in Supplementary Figure S16A-C. Most known disease genes display a significant “Gene Effect” on leukemia cell proliferation, while a few have not yet shown a measurable impact in the available cell line datasets. These findings are consistent with the results of our analysis of the predicted candidate genes. This further confirms the potential relevance of our predicted candidate genes to leukemia and validates the effectiveness of our prediction method.

**Figure 8.**
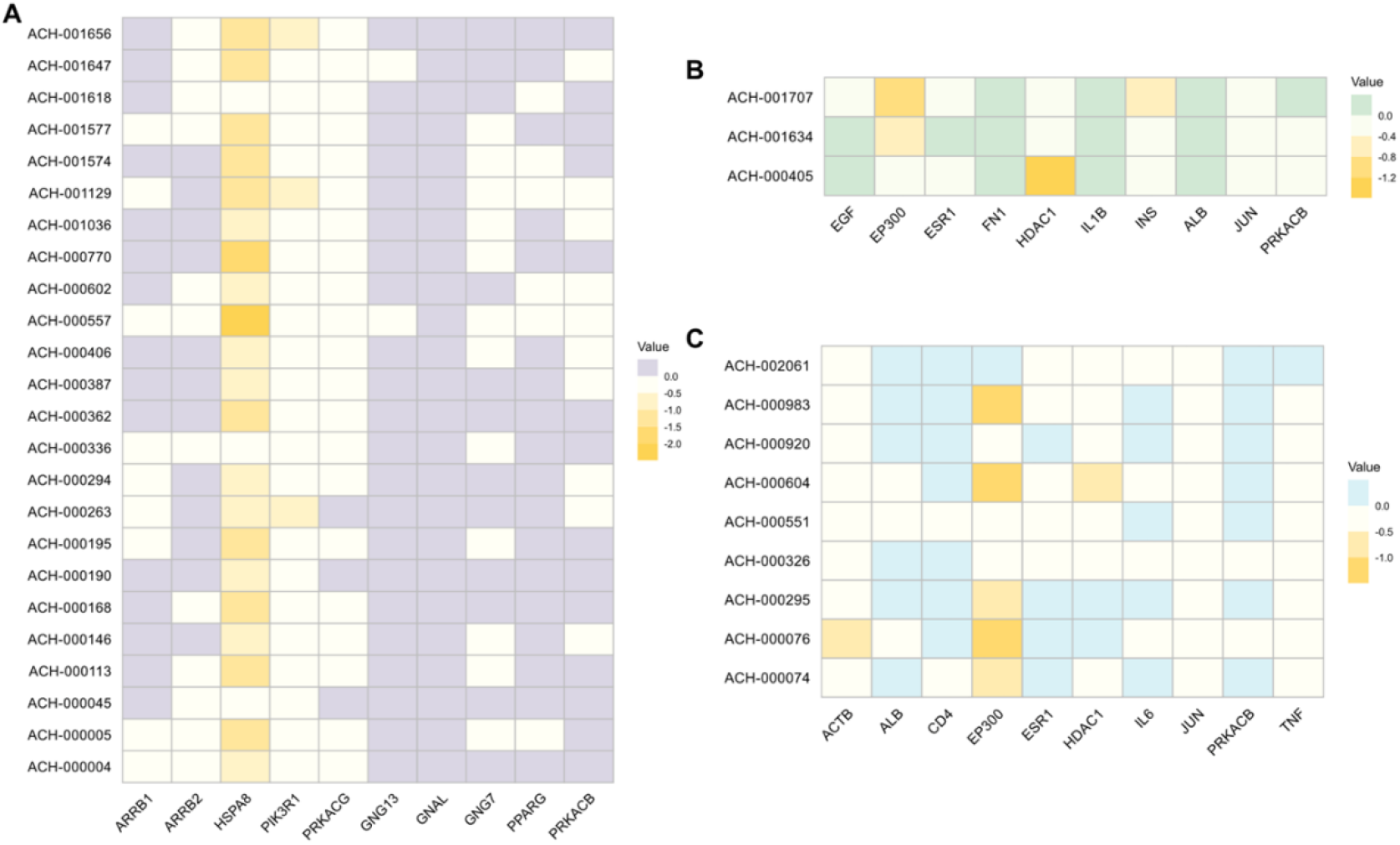
The “Gene Effect” values of the top ten predicted candidate genes for three types of leukemia. The heat map shows the “Gene Effect” values in corresponding cell lines for the top ten candidate genes predicted by DyNDG for three types of leukemia. **A**. Cell lines of AML. **B**. Cell lines of CLL. **C**. Cell lines of CML.

## Conclusion

The occurrence and development of blood diseases, such as leukemia, are closely related to gene mutations and abnormal expression. While existing methods for predicting disease genes have made significant contributions, most of them rely on static networks, limiting the improvement of their predictive ability. Addressing how to integrate dynamic information during disease development is a crucial issue that needs to be solved. We proposed the dynamic network-based model called uyNuG for predicting leukemia-related genes, which consists of three main steps: (1) construction of the time-series dynamic network; (2) construction of background-temporal multilayer biological network; and (3) the novel network propagation on the background-temporal multilayer network is proposed to obtain the ranking list of leukemia-related genes.

The uyNuG model was applied to the CLL-, CML-, and AML-related datasets for experiments. We evaluated the model performance in terms of four metrics: Topk_Precision, Topk_Recall, AUROC, and AUPRC. By comparing with other classical single-network, Multi-network methods, and methods using differential gene expression levels, it is found that our uyNuG model shows advantages in predicting leukemia-related genes.

Finally, we processed the candidate genes that uyNuG predicted for AML, CLL, and CML, separately, by excluding known leukemia-related genes and housekeeping genes, to generate three lists of candidate genes for the three types of leukemia. We systematically analyzed the candidate genes for the three types of leukemia from different perspectives by integrating various analysis methods. The results shown that the predicted candidate genes possess significant research value from various biological perspectives, further emphasizing the comprehensive predictive capability of our model, demonstrating the multifunctionality and robustness of our model accurately identifying disease genes across multiple types of leukemia.

## Data availability

The web server of DyNDG can be freely accessed at https://csuligroup.com/DyNDG

## Code availability

The implementation of DyNDG is available at https://github.com/CSUBioGroup/DyNDG.

## CRediT author statement

**Jin A**: Conceptualization, Data curation, Formal analysis, Methodology, Visualization, Writing - original draft

**Min Li** Formal analysis, Writing - review & editing, Supervision **Ju Xiang**: Formal analysis, Methodology, Writing original draft **Xiangmao Meng**: Data curation, Formal analysis

**Yue Sheng**: Writing - review & editing, Supervision

**Hongling Peng**: Writing - review & editing, Supervision

## Competing interests

The authors declare that they have no competing interests.

## Acknowledgements

This work was supported by grants from the National Natural Science Foundation of China (Grants No.62225209) [M.L.] and the National Natural Science Foundation of China (Grants No. 62472051) [J.X.].

## Notes

### Competing Interest Statement

The authors have declared no competing interest.

## References

[1] Sung H, Ferlay J, Siegel RL, Laversanne M, Soerjomataram I, Jemal A, et al. Global cancer statistics 2020: GLOBOCAN estimates of incidence and mortality worldwide for 36 cancers in 185 countries. CA Cancer J Clin 2021;71:209–49.

[2] Puckett Y, Chan O. Acute lymphocytic leukemia. In: StatPearls [Internet]. Treasure Island (FL): StatPearls Publishing 2023;Jan-Aug 26.

[3] Chang A, Schulz PJ, Wenghin Cheong A. Online newspaper framing of non-communicable diseases: comparison of Mainland China, Taiwan, Hong Kong and Macao. Int J Environ Res Public Health 2020;17:5593.

[4] Göring HH, Terwilliger JD, Blangero J. Large upward bias in estimation of locus-specific effects from genomewide scans. Am J Hum Genet 2001;69:1357–69.

[5] Lander E, Kruglyak L. Genetic dissection of complex traits: guidelines for interpreting and reporting linkage results. Nat Genet 1995;11:241–7.

[6] Sabatti C, Service S, Freimer N. False discovery rate in linkage and association genome screens for complex disorders. Genetics 2003;164:829–33.

[7] Luo P, Tian L-P, Ruan J, Wu F-X. Disease gene prediction by integrating ppi networks, clinical rna-seq data and omim data. IEEE/ACM Trans Comput Biol Bioinform 2017;16:222–32.

[8] Barabási A-L, Gulbahce N, Loscalzo J. Network medicine: a network-based approach to human disease. Nat Rev Genet 2011;12:56–68.

[9] Yue X, Wang Z, Huang J, Parthasarathy S, Moosavinasab S, Huang Y, et al. Graph embedding on biomedical networks: methods, applications and evaluations. Bioinformatics 2020;36:1241–51.

[10] Wu X, Jiang R, Zhang MQ, Li S. Network-based global inference of human disease genes. Mol Syst Biol 2008;4:189.

[11] Sun PG, Gao L, Han S. Prediction of human disease-related gene clusters by clustering analysis. Int J Biol Sci 2011;7:61.

[12] Oti M, Snel B, Huynen MA, Brunner HG. Predicting disease genes using protein–protein interactions. J Med Genet 2006;43:691–8.

[13] Lage K, Karlberg EO, Størling ZM, Olason PI, Pedersen AG, Rigina O, et al. A human phenome-interactome network of protein complexes implicated in genetic disorders. Nat Biotechnol 2007;25:309–16.

[14] Ata SK, Wu M, Fang Y, Ou-Yang L, Kwoh CK, Li X-L. Recent advances in network-based methods for disease gene prediction. Brief Bioinform 2021;22:bbaa303.

[15] Hu K, Hu J-B, Tang L, Xiang J, Ma J-L, Gao Y-Y, et al. Predicting disease-related genes by path structure and community structure in protein–protein networks. J Stat Mech: Theory Exp 2018;2018:100001.

[16] Xiang J, Meng X, Zhao Y, Wu F-X, Li M. HyMM: hybrid method for disease-gene prediction by integrating multiscale module structure. Brief Bioinform 2022;23:bbac072.

[17] Li Y, Patra JC. Genome-wide inferring gene–phenotype relationship by walking on the heterogeneous network. Bioinformatics 2010;26:1219–24.

[18] Li Y, Li J. Disease gene identification by random walk on multigraphs merging heterogeneous genomic and phenotype data. BMC Genomics 2012;13:1–12.

[19] Köhler S, Bauer S, Horn D, Robinson PN. Walking the interactome for prioritization of candidate disease genes. Am J Hum Genet 2008;82:949–58.

[20] Valdeolivas A, Tichit L, Navarro C, Perrin S, Odelin G, Levy N, et al. Random walk with restart on multiplex and heterogeneous biological networks. Bioinformatics 2019;35:497–505.

[21] Zhang Y, Xiang J, Tang L, Li J, Lu Q, Tian G, et al. Identifying breast cancer-related genes based on a novel computational framework involving KEGG pathways and PPI network modularity. Front Genet 2021;12:596794.

[22] Przytycka TM, Singh M, Slonim DK. Toward the dynamic interactome: it’s about time. Brief Bioinform 2010;11:15–29.

[23] Hegele A, Kamburov A, Grossmann A, Sourlis C, Wowro S, Weimann M, et al. Dynamic protein-protein interaction wiring of the human spliceosome. Mol Cell 2012;45:567–80.

[24] Carneiro DG, Clarke T, Davies CC, Bailey D. Identifying novel protein interactions: Proteomic methods, optimisation approaches and data analysis pipelines. Methods 2016;95:46–54.

[25] Wang J, Peng X, Peng W, Wu FX. Dynamic protein interaction network construction and applications. Proteomics 2014;14:338–52.

[26] de Lichtenberg U, Jensen LJ, Brunak S, Bork P. Dynamic complex formation during the yeast cell cycle. Science 2005;307:724–7.

[27] Hegde SR, Manimaran P, Mande SC. Dynamic changes in protein functional linkage networks revealed by integration with gene expression data. PLoS Comput Biol 2008;4:e1000237.

[28] Tang X, Wang J, Liu B, Li M, Chen G, Pan Y. A comparison of the functional modules identified from time course and static PPI network data. BMC Bioinformatics 2011;12:1–15.

[29] Zhang Y, Lin H, Yang Z, Wang J. Construction of dynamic probabilistic protein interaction networks for protein complex identification. BMC Bioinformatics 2016;17:1–13.

[30] Ou-Yang L, Dai D-Q, Li X-L, Wu M, Zhang X-F, Yang P. Detecting temporal protein complexes from dynamic protein-protein interaction networks. BMC Bioinformatics 2014;15:1–14.

[31] Lei X, Wang F, Wu F-X, Zhang A, Pedrycz W. Protein complex identification through Markov clustering with firefly algorithm on dynamic protein–protein interaction networks. Inform Sciences 2016;329:303–16.

[32] Xiao Q, Wang J, Peng X, Wu F-x, Pan Y. Identifying essential proteins from active PPI networks constructed with dynamic gene expression. BMC Genomics 2015;16:1–7.

[33] Li M, Chen X, Ni P, Wang J, Pan Y. Identifying essential proteins by purifying protein interaction networks. ISBRA 2016:106–16.

[34] Chen L, Liu R, Liu Z-P, Li M, Aihara K. Detecting early-warning signals for sudden deterioration of complex diseases by dynamical network biomarkers. Sci Rep 2012;2:1–8.

[35] Liu R, Wang X, Aihara K, Chen L. Early diagnosis of complex diseases by molecular biomarkers, network biomarkers, and dynamical network biomarkers. Med Res Rev 2014;34:455–78.

[36] Taylor IW, Linding R, Warde-Farley D, Liu Y, Pesquita C, Faria D, et al. Dynamic modularity in protein interaction networks predicts breast cancer outcome. Nat Biotechnol 2009;27:199–204.

[37] Faisal FE, Milenkovic T. Dynamic networks reveal key players in aging. Bioinformatics 2014;30:1721–9.

[38] Chennamadhavuni A, Lyengar V, Shimanovsky A. Leukemia. In: StatPearls [Internet]. Treasure Island (FL): StatPearls Publishing 2022;Jan 17.

[39] Board PATE. Acute Myeloid Leukemia Treatment (PDQ®). PDQ Cancer Information Summaries [Internet]. National Cancer Institute (US), 2022.

[40] Board PATE. Chronic Lymphocytic Leukemia Treatment (PDQ®). PDQ Cancer Information Summaries [Internet]. National Cancer Institute (US), 2022.

[41] Board PATE. Chronic Myelogenous Leukemia Treatment (PDQ®). PDQ Cancer Information Summaries [Internet]. National Cancer Institute (US), 2022.

[42] Szklarczyk D, Kirsch R, Koutrouli M, Nastou K, Mehryary F, Hachilif R, et al. The STRING database in 2023: protein-protein association networks and functional enrichment analyses for any sequenced genome of interest. Nucleic Acids Res 2023;51(D1):D638–D646.

[43] Kim CY, Baek S, Cha J, Yang S, Kim E, Marcotte EM, et al. HumanNet v3: an improved database of human gene networks for disease research. Nucleic Acids Res 2022;50(D1):D632–D9.

[44] Rappaport N, Twik M, Plaschkes I, Nudel R, Iny Stein T, Levitt J, et al. MalaCards: an amalgamated human disease compendium with diverse clinical and genetic annotation and structured search. Nucleic Acids Res 2017;45(D1):D877–D87.

[45] Seal RL, Braschi B, Gray K, Jones TE, Tweedie S, Haim-Vilmovsky L, et al. Genenames. org: the HGNC resources in 2023. Nucleic Acids Res 2023;51(D1):D1003–D1009.

[46] Li Y, Patra JC. Integration of multiple data sources to prioritize candidate genes using discounted rating system. BMC Bioinformatics 2010;11:1–10.

[47] Ruan P, Wang S. DiSNEP: a disease-specific gene network enhancement to improve prioritizing candidate disease genes. Brief Bioinform 2021;22:bbaa241.

[48] Hounkpe BW, Chenou F, de Lima F, De Paula EV. HRT Atlas v1. 0 database: redefining human and mouse housekeeping genes and candidate reference transcripts by mining massive RNA-seq datasets. Nucleic Acids Res 2021;49(D1):D947–D55.

[49] Li K, Du Y, Cai Y, Liu W, Lv Y, Huang B, et al. Single-cell analysis reveals the chemotherapy-induced cellular reprogramming and novel therapeutic targets in relapsed/refractory acute myeloid leukemia. Leukemia 2023;37:308–25.

[50] Bruserud Ø, Reikvam H, Brenner AK. Toll-like receptor 4, osteoblasts and leukemogenesis; the lesson from acute myeloid leukemia. Molecules 2022;27:735.

[51] He M, Jia Y, Wang Y, Cai X. Dysregulated MAPK signaling pathway in acute myeloid leukemia with RUNX1 mutations. Expert Rev Hematol. 2022;15(8):769–779.

[52] Hsiao P-C, Hsieh Y-H, Chow J-M, Yang S-F, Hsiao M, Hua K-T, et al. Hispolon induces apoptosis through JNK1/2-mediated activation of a caspase-8,-9, and-3-dependent pathway in acute myeloid leukemia (AML) cells and inhibits AML xenograft tumor growth in vivo. J Agric Food Chem 2013;61:10063–73.

[53] Su Y-C, Li S-C, Wu Y-C, Wang L-M, Chao K, Liao H-F. Resveratrol downregulates interleukin-6-stimulated sonic hedgehog signaling in human acute myeloid leukemia. Evid Based Complement Alternat Med 2013;2013.

[54] Lu FL, Yu C-C, Chiu H-H, Liu HE, Chen S-Y, Lin S, et al. Sonic hedgehog antagonists induce cell death in acute myeloid leukemia cells with the presence of lipopolysaccharides, tumor necrosis factor-α, or interferons. Invest New Drugs 2013;31:823–32.

[55] Ibanez M, Carbonell-Caballero J, Such E, Garcia-Alonso L, Liquori A, Lopez-Pavia M, et al. The modular network structure of the mutational landscape of Acute Myeloid Leukemia. PLoS One 2018;13:e0202926.

[56] Darici S, Alkhaldi H, Horne G, Jørgensen HG, Marmiroli S, Huang X. Targeting PI3K/Akt/mTOR in AML: rationale and clinical evidence. J Clin Med 2020;9:2934.

[57] Patel K, Pagel JM. Current and future treatment strategies in chronic lymphocytic leukemia. J Hematol Oncol 2021;14:1–20.

[58] Shannon P, Markiel A, Ozier O, Baliga NS, Wang JT, Ramage D, et al. Cytoscape: a software environment for integrated models of biomolecular interaction networks. Genome Res 2003;13:2498–504.

[59] Wang J, Kafeel M, Avezbakiyev B, Chen C, Sun Y, Rathnasabapathy C, et al. Histone deacetylase in chronic lymphocytic leukemia. Oncology 2012;81:325–9.

[60] Lai T-H, Ozer HG, Gasparini P, Nigita G, Distefano R, Yu L, et al. HDAC1 regulates the chromatin landscape to control transcriptional dependencies in chronic lymphocytic leukemia. Blood Adv 2023;7:2897–911.

[61] Byrd JC, Woyach JA, Johnson AJ. Translating PI3K-delta inhibitors to the clinic in chronic lymphocytic leukemia: the story of CAL-101 (GS1101). Am Soc Clin Oncol Educ Book 2012;32:691–4.

[62] Durand-Onayli V, Haslauer T, Härzschel A, Hartmann TN. Rac GTPases in hematological malignancies. Int J Mol Sci 2018;19:4041.

[63] Shaffer III AL, Phelan JD, Wang JQ, Huang D, Wright GW, Kasbekar M, et al. Overcoming acquired epigenetic resistance to BTK inhibitors. Blood Cancer Discov 2021;2:630–47.

[64] Minciacchi VR, Kumar R, Krause DS. Chronic myeloid leukemia: a model disease of the past, present and future. Cells 2021;10:117.

[65] Lee CR, Kang JA, Kim HE, Choi Y, Yang T, Park SG. Secretion of IL-1β from imatinib-resistant chronic myeloid leukemia cells contributes to BCR-ABL mutation-independent imatinib resistance. FEBS Lett 2016;590:358–68.

[66] Bütow M, Testaquadra FJ, Baumeister J, Maié T, Chatain N, Jaquet T, et al. Targeting cytokine-induced leukemic stem cell persistence in chronic myeloid leukemia by IKK2-inhibition. Haematologica 2023;108:1179.

[67] Sharma K, Singh U, Rai M, Shukla J, Gupta V, Narayan G, et al. Interleukin 6 and disease transformation in chronic myeloid leukemia: A Northeast Indian population study. J Cancer Res Ther 2020;16:30–3.

[68] Li M, ZHOU L-j. Research progress on the relationship between TRAF6 and tumor. Biotechnology Bulletin 2017;33:24.

[69] Han SH, Korm S, Han YG, Choi S-Y, Kim S-H, Chung HJ, et al. GCA links TRAF6-ULK1-dependent autophagy activation in resistant chronic myeloid leukemia. Autophagy 2019;15:2076–90.

[70] Mencalha AL, Corrêa S, Abdelhay E. Role of calcium-dependent protein kinases in chronic myeloid leukemia: combined effects of PKC and BCR-ABL signaling on cellular alterations during leukemia development. OncoTargets Ther 2014:1247–54.

[71] Tsherniak A, Vazquez F, Montgomery PG, Weir BA, Kryukov G, Cowley GS, et al. Defining a cancer dependency map. Cell 2017;170:564-76. e16.

[72] Li J, Ge Z. High HSPA8 expression predicts adverse outcomes of acute myeloid leukemia. BMC Cancer 2021;21:1–11.

[73] Zhu Y, Wang Z, Li Y, Peng H, Liu J, Zhang J, et al. The Role of CREBBP/EP300 and Its Therapeutic Implications in Hematological Malignancies. Cancers 2023;15:1219.

