## Supplementary material for "DyNDG: Identifying Leukemia-Related Genes based on the Time-Series Dynamical Network by Integrating Differential Genes": File S1.docx

**Section 1. Calculation process of Formula (5)**

Let $W_{m}$ be a $N\times N$ weighting matrix obtained by replicating the vector ${{AP}_{m}}^{T}$ $N$ times. The reweighted adjacency matrix $A_{m}^{'}$ for patient sample $m$ is calculated as follows:

$$\begin{aligned} A_{m}^{'} ={{(A}_{m}\odot W_{m})}^{T}\odot W_{m} \end{aligned}$$

$$\begin{aligned} W_{m} = \left[ {{AP}_{m}}^{T},{{AP}_{m}}^{T},\ldots,{{AP}_{m}}^{T} \right] \in R^{N\times N} \end{aligned}$$

**Section 2. Example and parameter description of Formula (9)**

For Formula (9), if there are 4 stages, S = 4 and

$$\begin{aligned} T_{S} = \left\{ \begin{matrix} \begin{matrix} \left( 1-\mu\right)A^{\left[ 1 \right]} & \mu I \\ \mu I & \left( 1-\mu\right)A^{\left[ 2 \right]} \end{matrix} & \begin{matrix} 0 & 0 \\ \mu I & 0 \end{matrix} \\ \begin{matrix} 0 & \mu I \\ 0 & 0 \end{matrix} & \begin{matrix} \left( 1-\mu\right)A^{\left[ 3 \right]} & \mu I \\ \mu I & \left( 1-\mu\right)A^{\left[ 4 \right]} \end{matrix} \end{matrix} \right\} \end{aligned}$$

When S=3 and 5, substitute the value of S into Formula (9) to get $T_{S}$. $\mu$ adjusts the transition between the time-series dynamic network layers. The meanings of $\mu=0, 0.5,$and $1$ are respectively:

1) $\mu=0$: Only intra-layer jumps, no direct transitions between the time-series dynamic network layers.

2) $\mu=0.5$: There are links with a weight of 0.5 between consecutive time-series dynamic network layers.

3) $\mu=1:$ Only inter-layer jumps, no intra-layer transitions.

**Section 3. Analysis of parameter setting**

We investigated the changes in the AUROC and AUPRC metrics of the DyNDG model concerning the values of parameter $\delta$, $\mu$, and $\gamma$ in three types of leukemia using the WG control set. The parameter $\delta$ adjusts the transition probability between the background network and the multi-layer dynamic network; $\mu$ adjusts the transition between the time-series dynamic network layers; and $\gamma$ adjusts the restart probability. Using grid search, we systematically studied the influence of the above parameters on the model performance. Analyzing the changes of AUROC and AUPRC, as shown in Supplementary Figure S6-S11, we observed no significant changes in AUROC and AUPRC for different values of the parameters $\delta$ (from $\delta=0.1$ to $\delta=0.8$) and $\mu$ (from $\mu=0.1$ to $\mu=1.0$). As $\gamma$ varies from 0.1 to 0.9, the AUROC and AUPRC values of the DyNDG model decrease. Therefore, we chose $\delta=0.5, \mu=0.5, \gamma=0.1$ as the parameter combination yields the relatively best model performance.

**Section 4. Effect of different static network datasets**

When different static networks are used, DyNDG may exhibit different predictive abilities. We explored the influence of different static networks (STRING and HumanNet) on the AUROC and AUPRC metrics using three control sets. As shown in Supplementary Figure S12, DyNDG performs better when using the static PPI network of STRING.

**Section 5. Experimental setup for ablation study**

For DyNDG_static,

$$T_{L} = B$$

The random-walk process is performed exclusively on the background network. It retrieves the score for each gene from the background network.

For DyNDG_dynet,

$$T_{L} = \left[ \begin{matrix} 0 & \delta J^{T} \\ \delta J & T_{S} \end{matrix} \right]$$

Set $\delta$ to 1, keeping the connections between the layers of the network, but the edge weights within the background network are set to 0. This is equivalent to having the random walk process occur only on the dynamic network, with the background network structure having no effect. The steady-state scoring vector $P_{\infty}^{B}$ on the background network is still used to evaluate the association between genes and leukemia.
