## Supplementary figures and images for "DyNDG: Identifying Leukemia-Related Genes based on the Time-Series Dynamical Network by Integrating Differential Genes"

### Figure S1.pptx

#### Slide 1
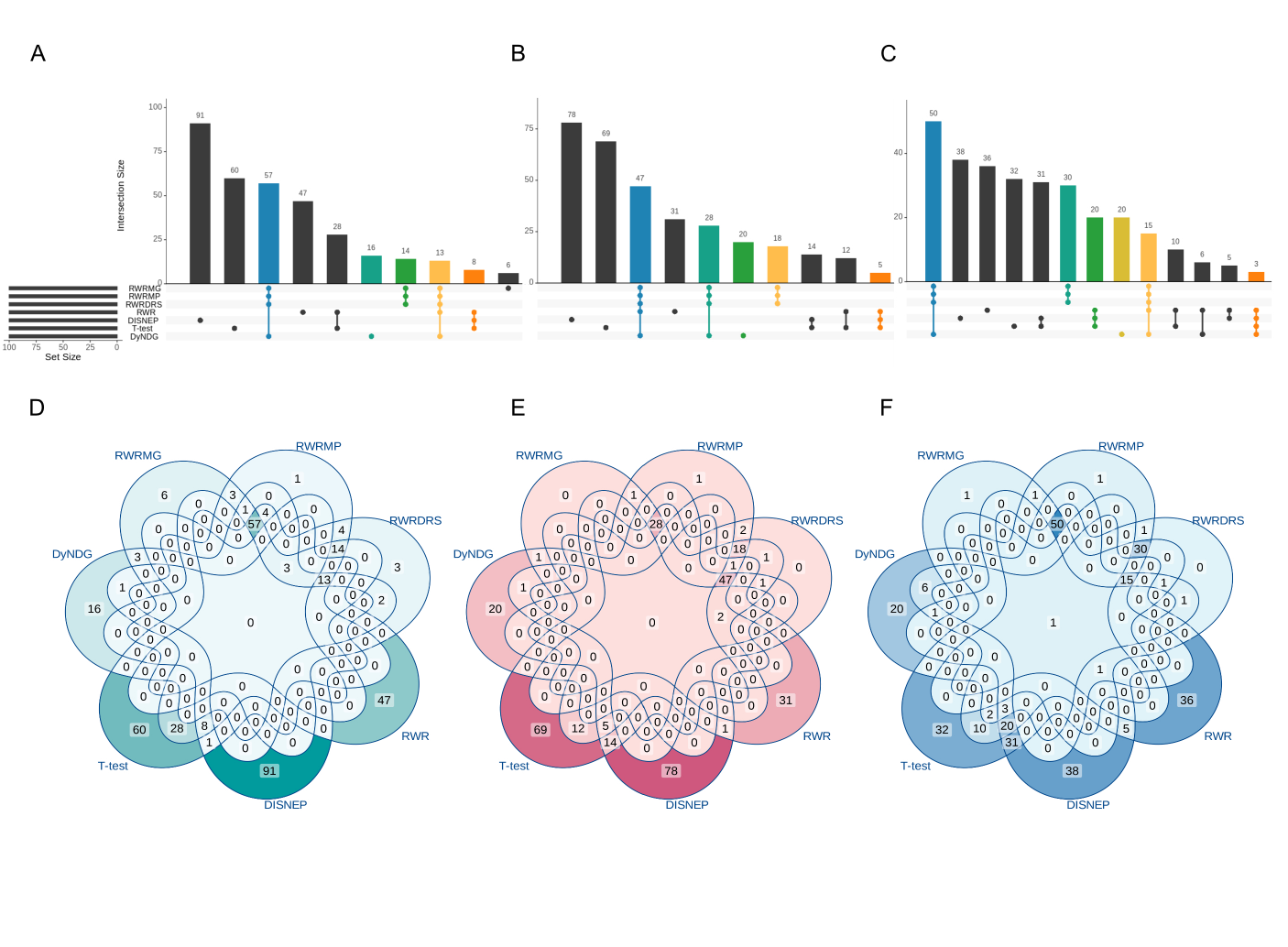

### Figure S2.pptx

#### Slide 1
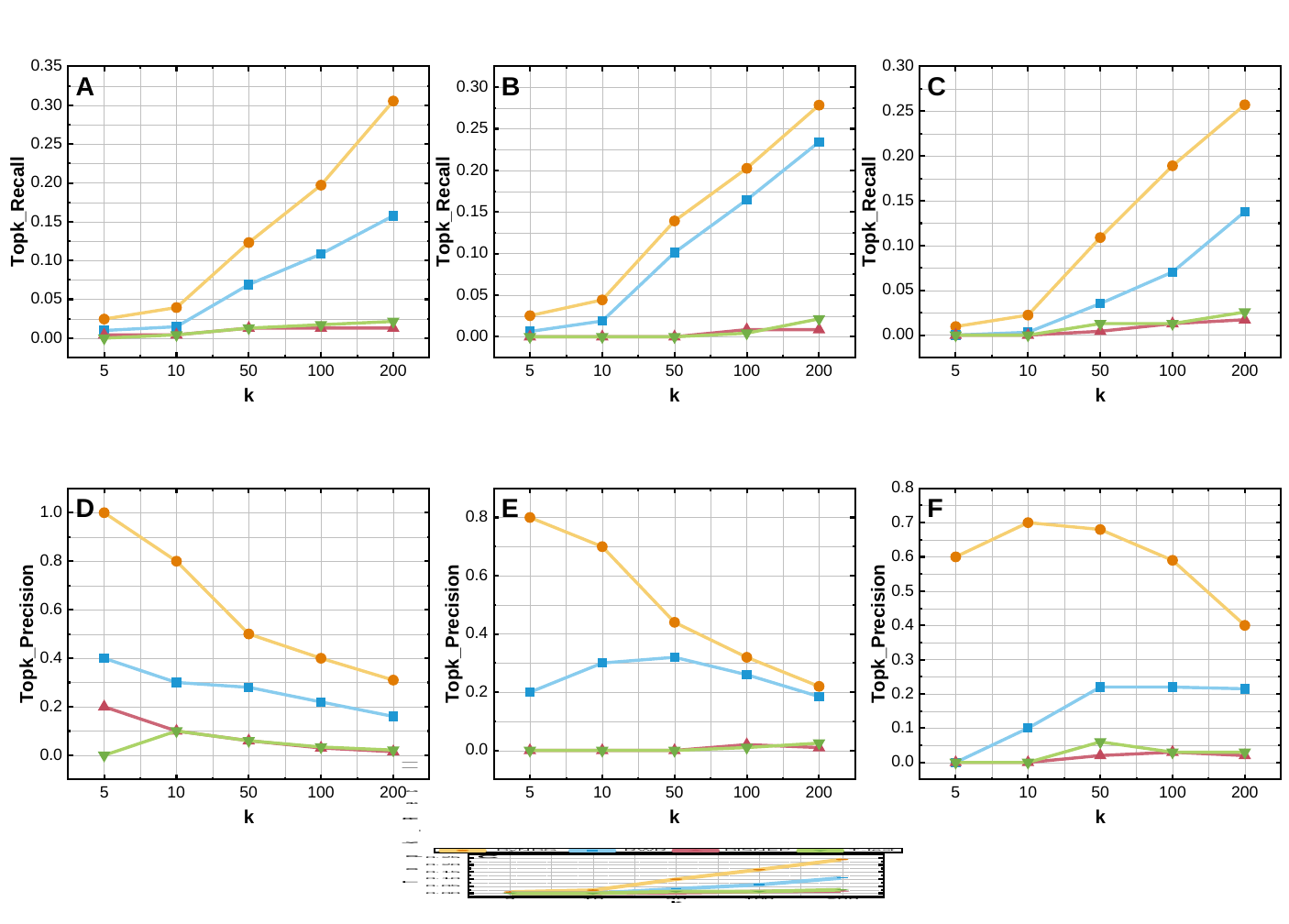

### Figure S3.pptx

#### Slide 1
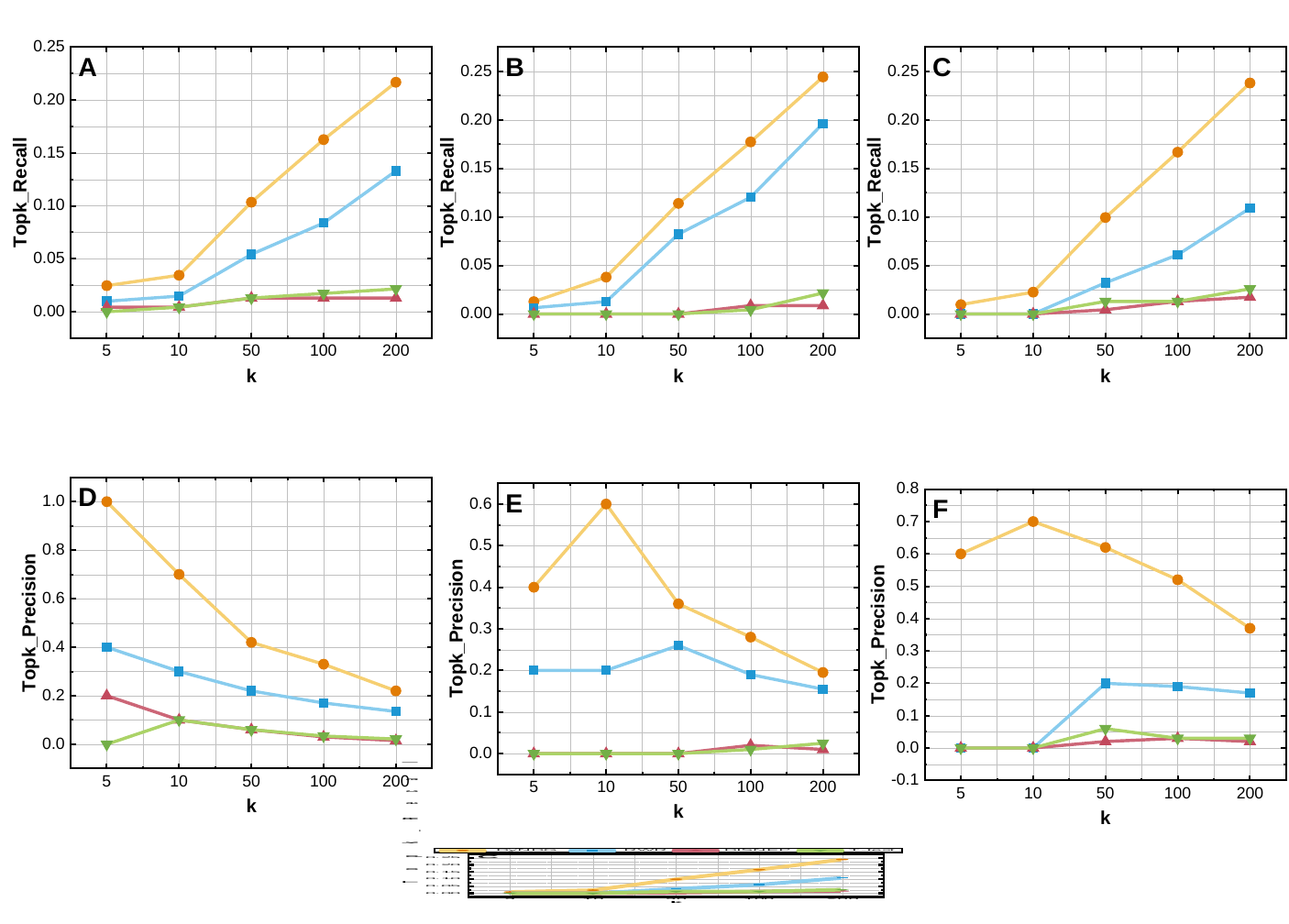

### Figure S4.pptx

#### Slide 1
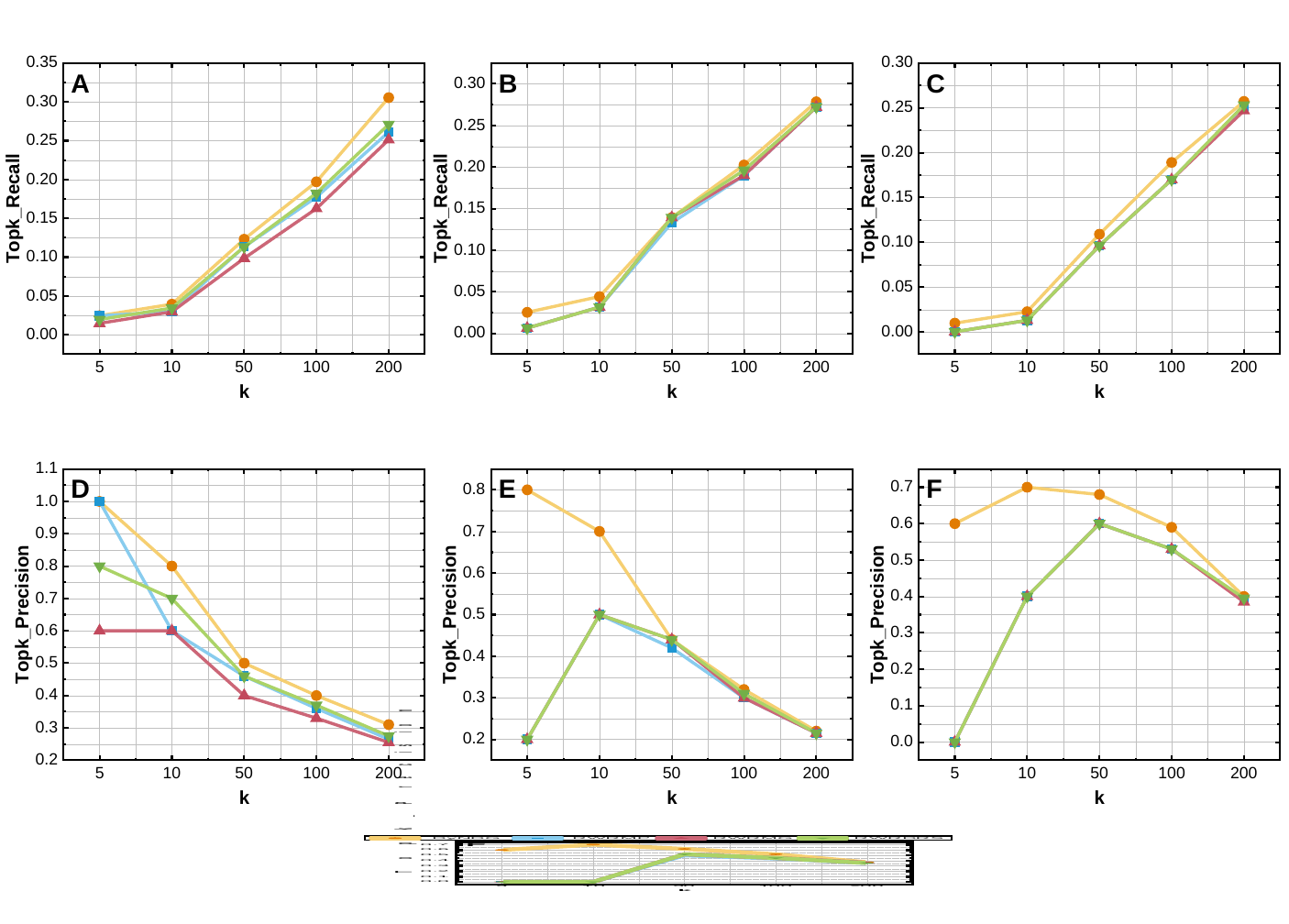

### Figure S5.pptx

#### Slide 1
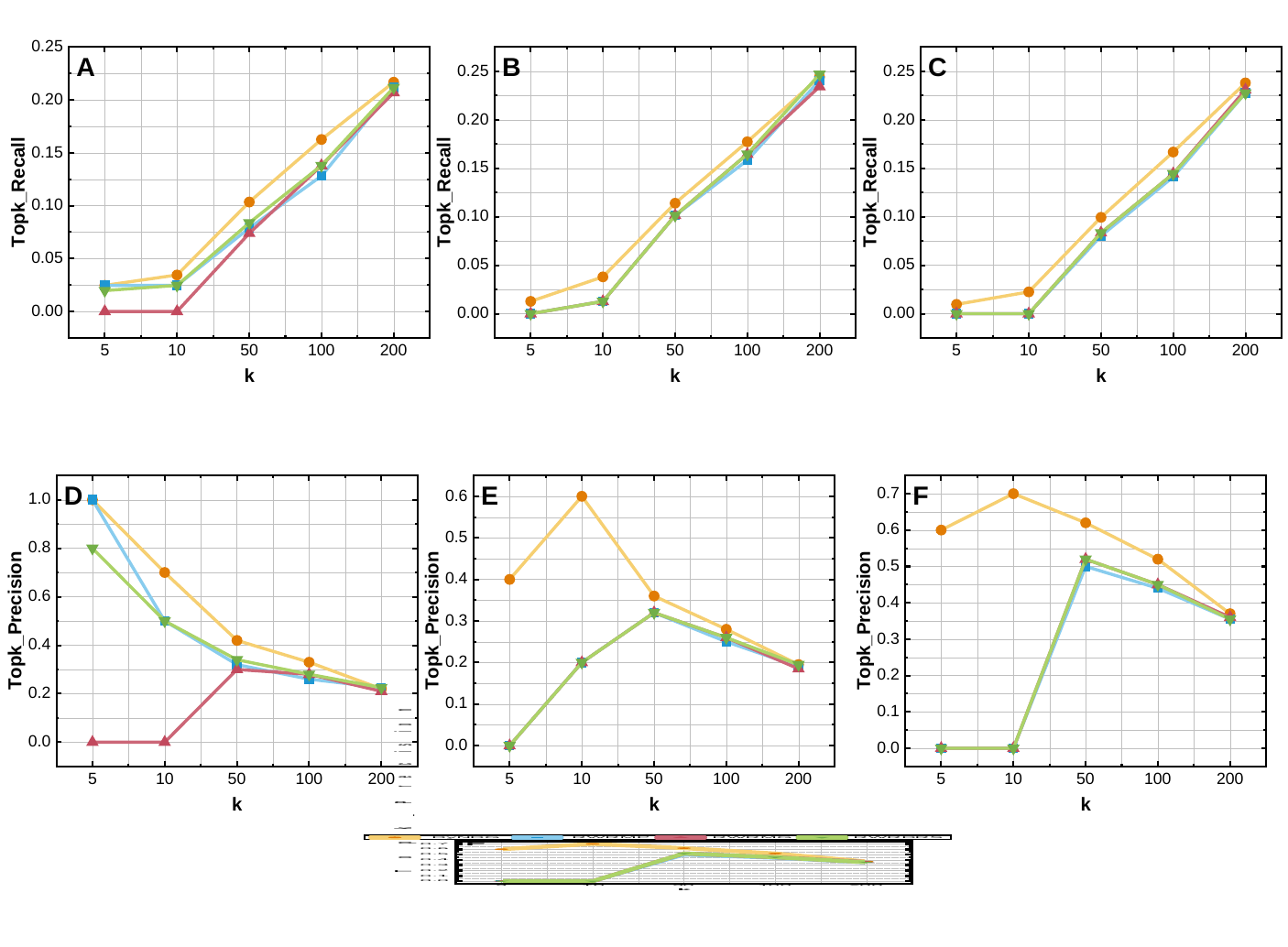

### Figure S6.pptx

#### Slide 1
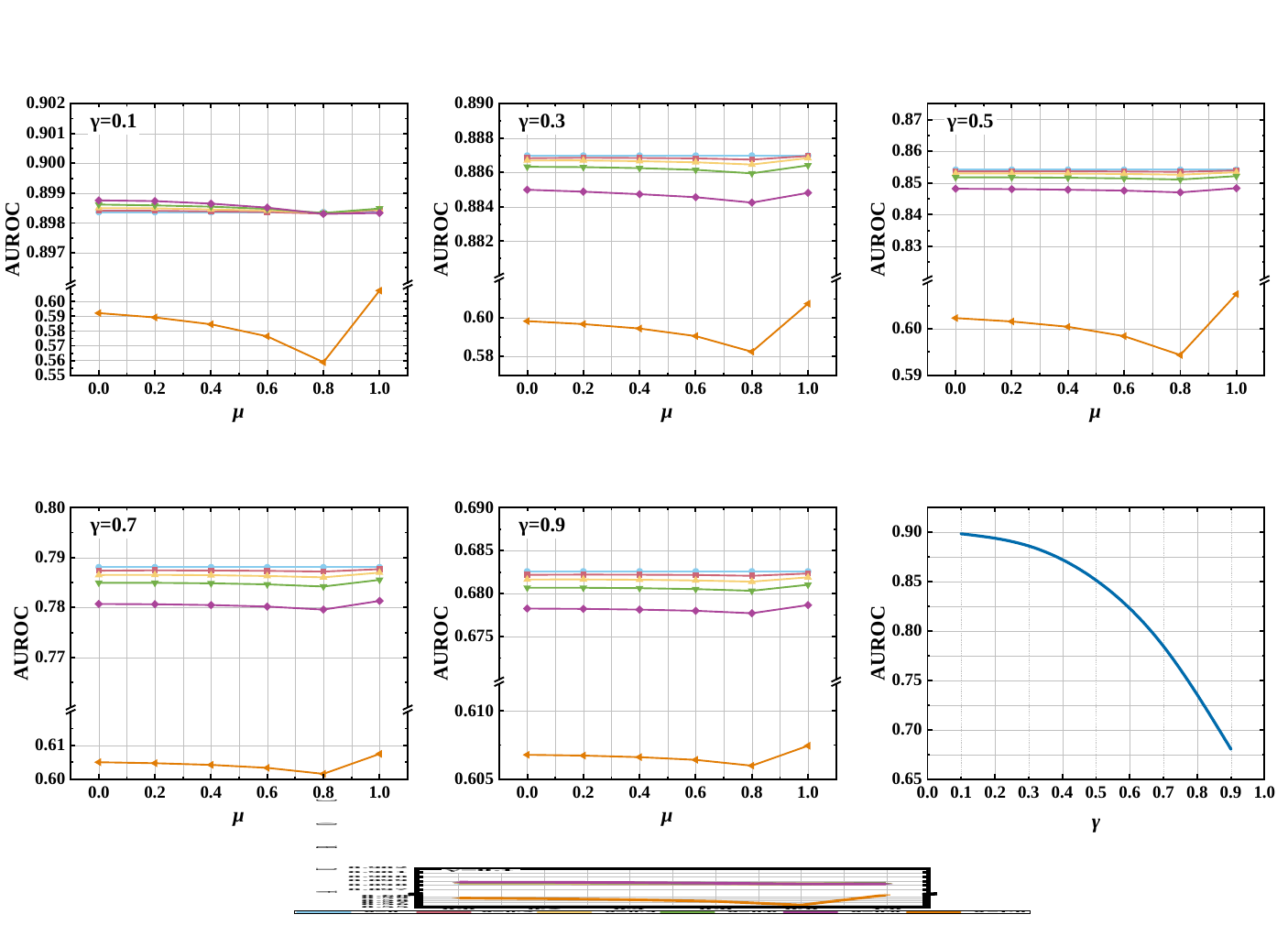

### Figure S7.pptx

#### Slide 1
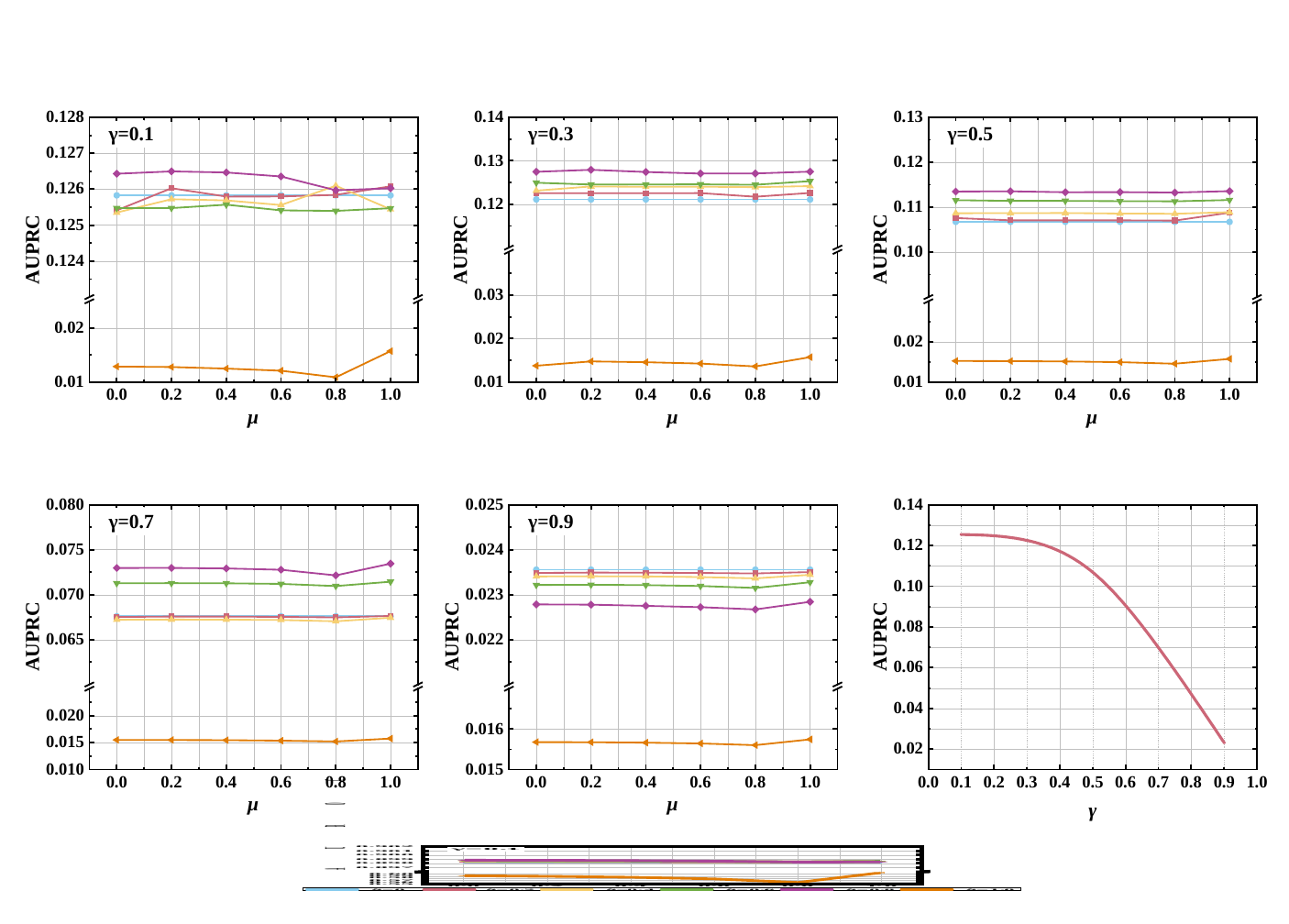

1

### Figure S8.pptx

#### Slide 1
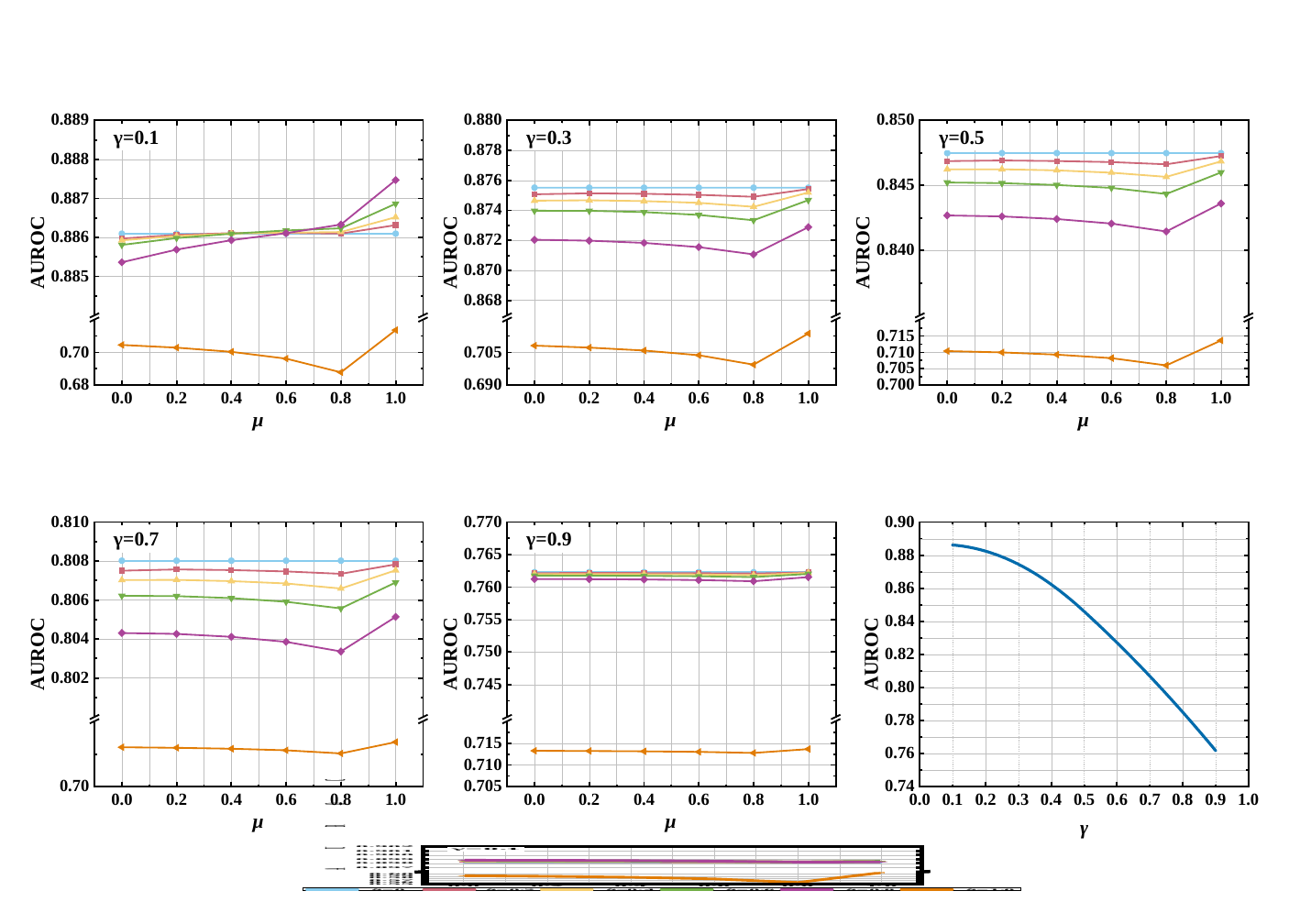

1

### Figure S9.pptx

#### Slide 1
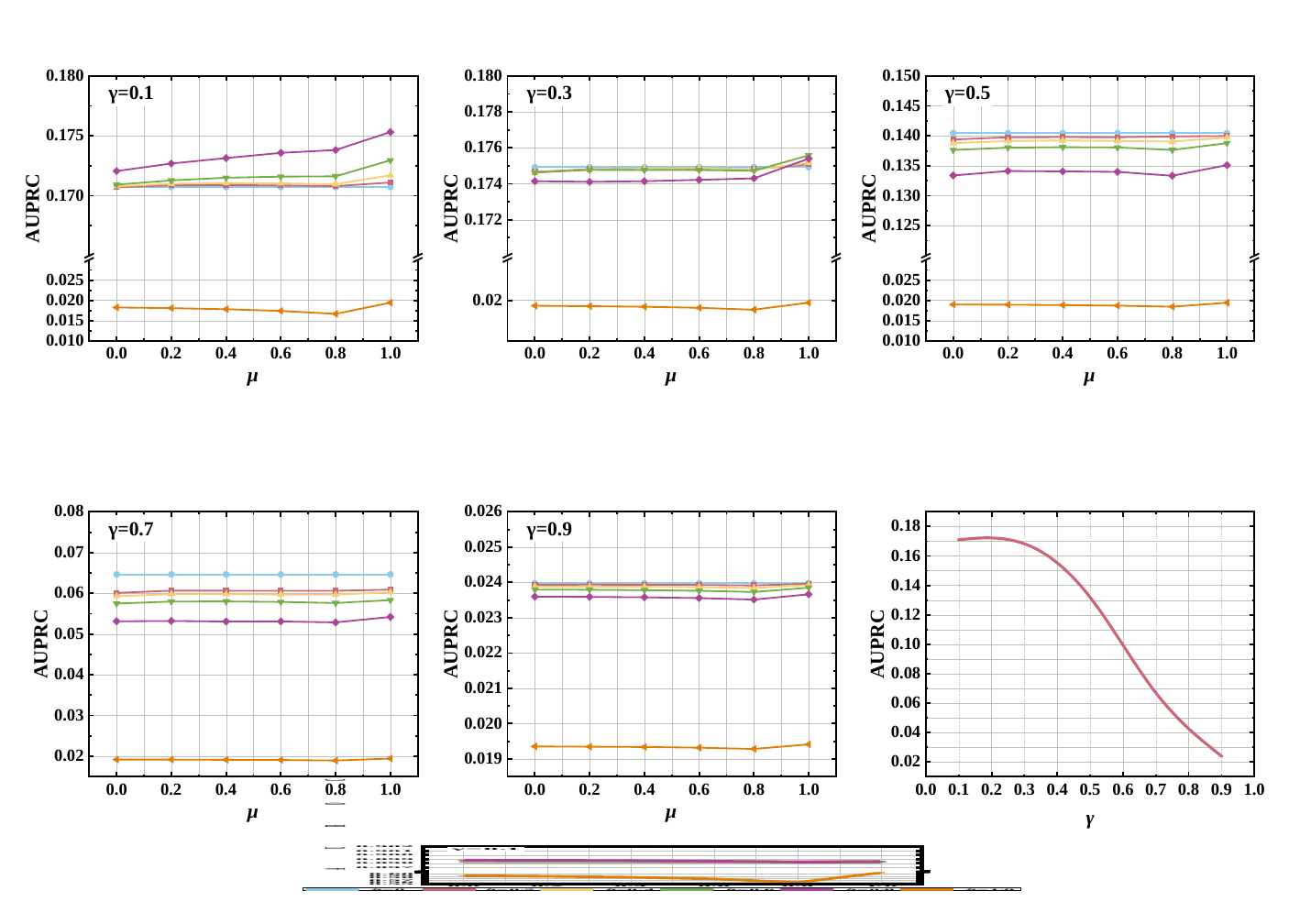

1

### Figure S10.pptx

#### Slide 1
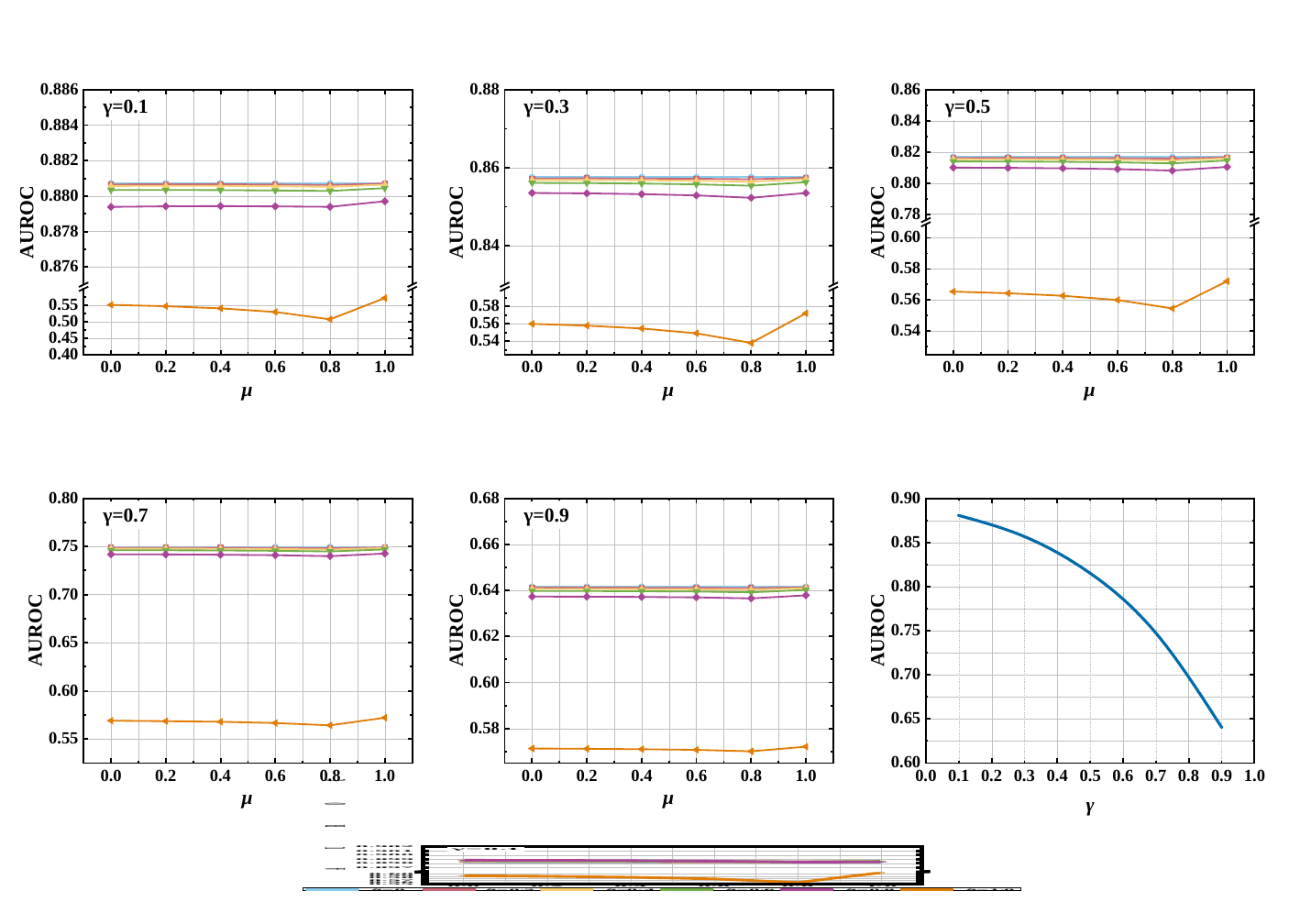

1

### Figure S11.pptx

#### Slide 1
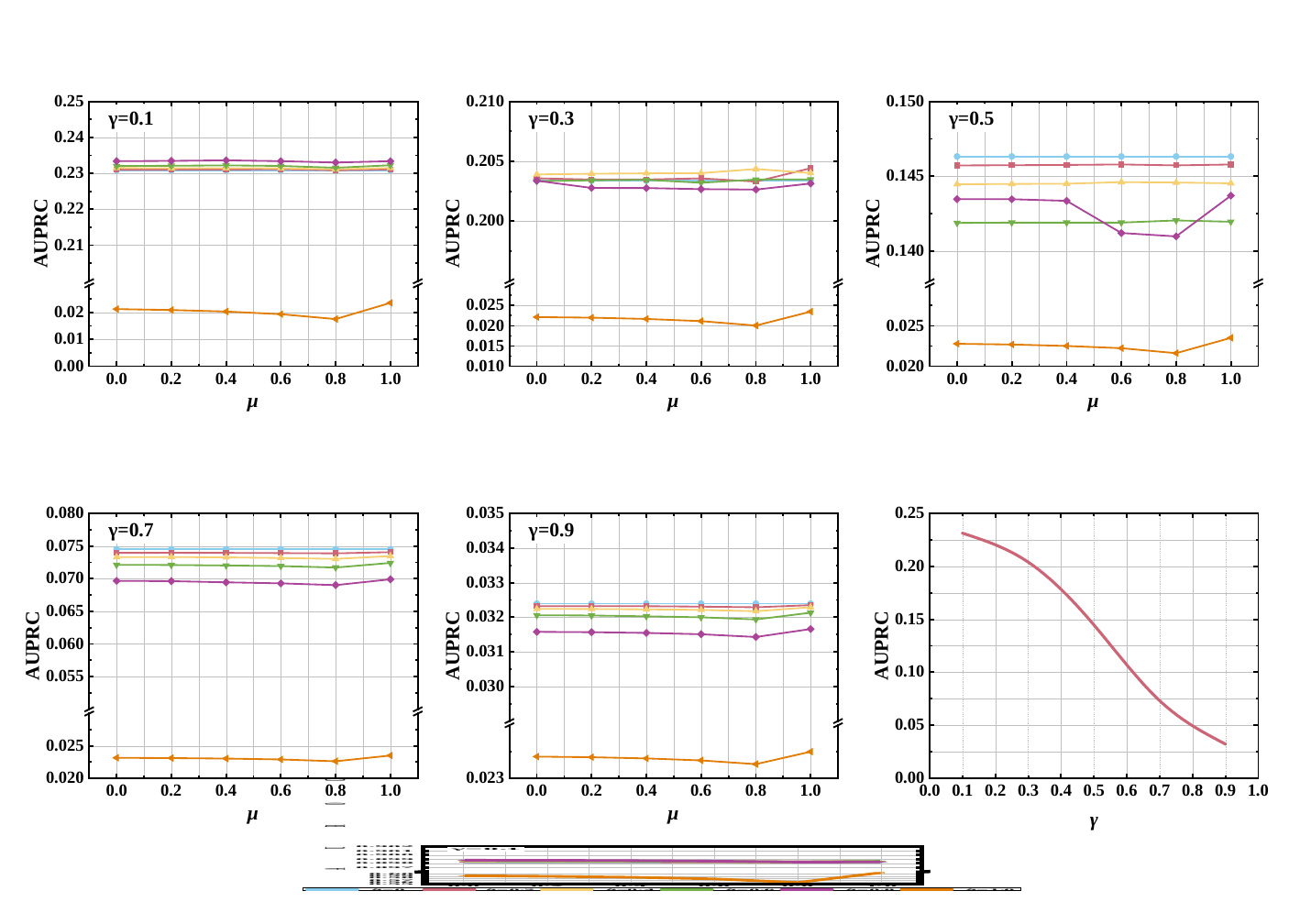

1

### Figure S12.pptx

#### Slide 1
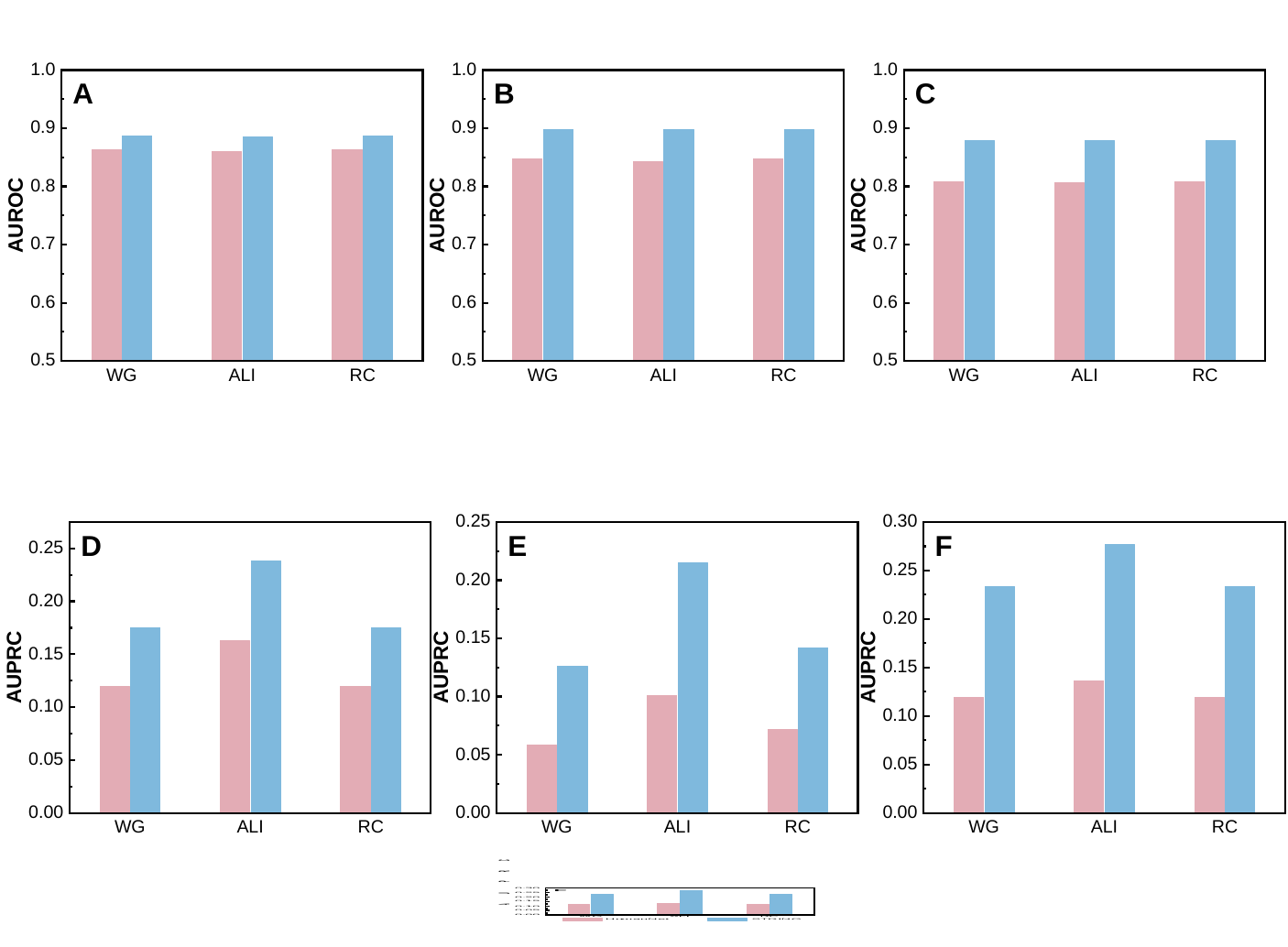

### Figure S13.pptx

#### Slide 1
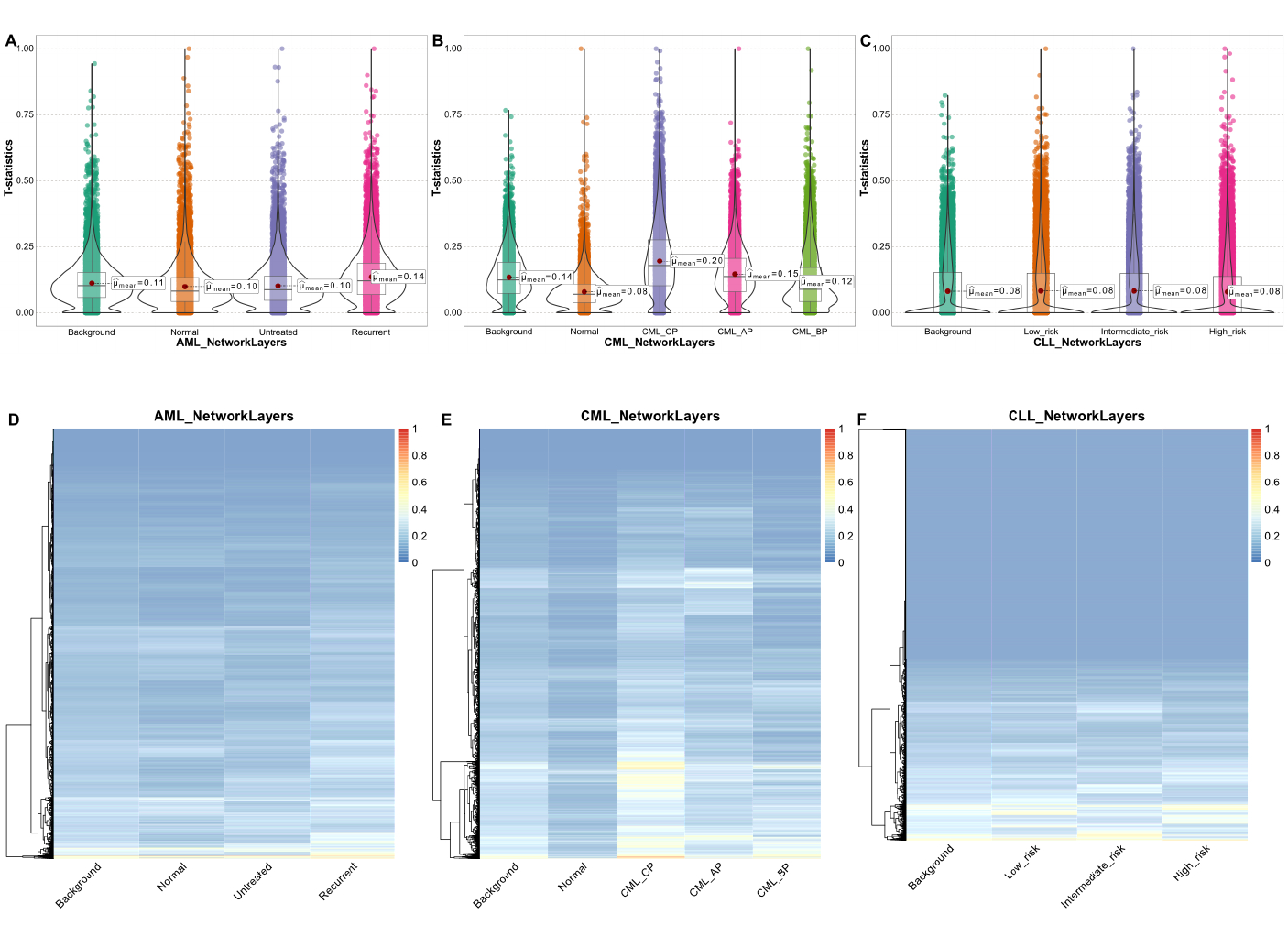

### Figure S14.pptx

#### Slide 1
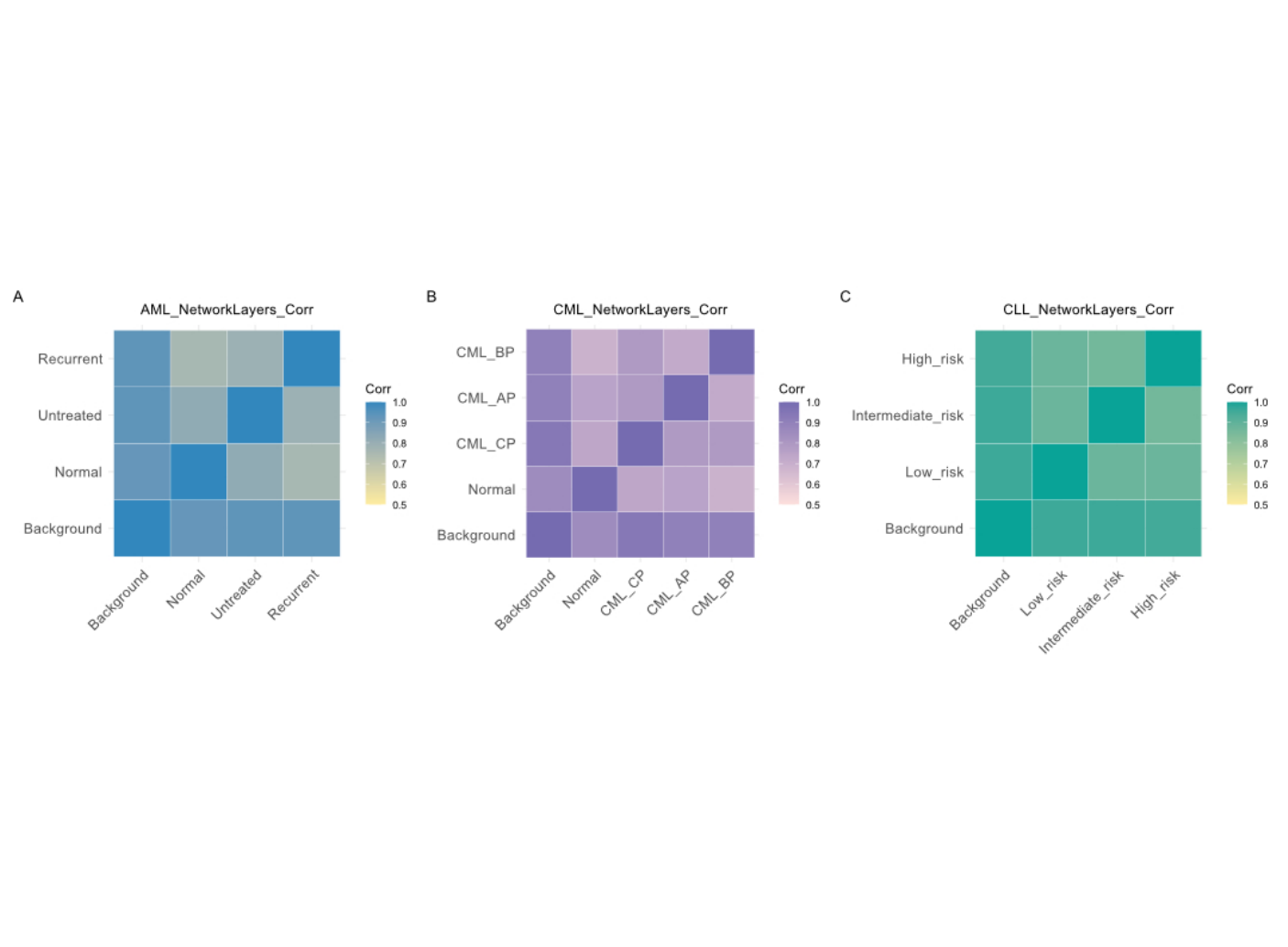

### Figure S15.pptx

#### Slide 1
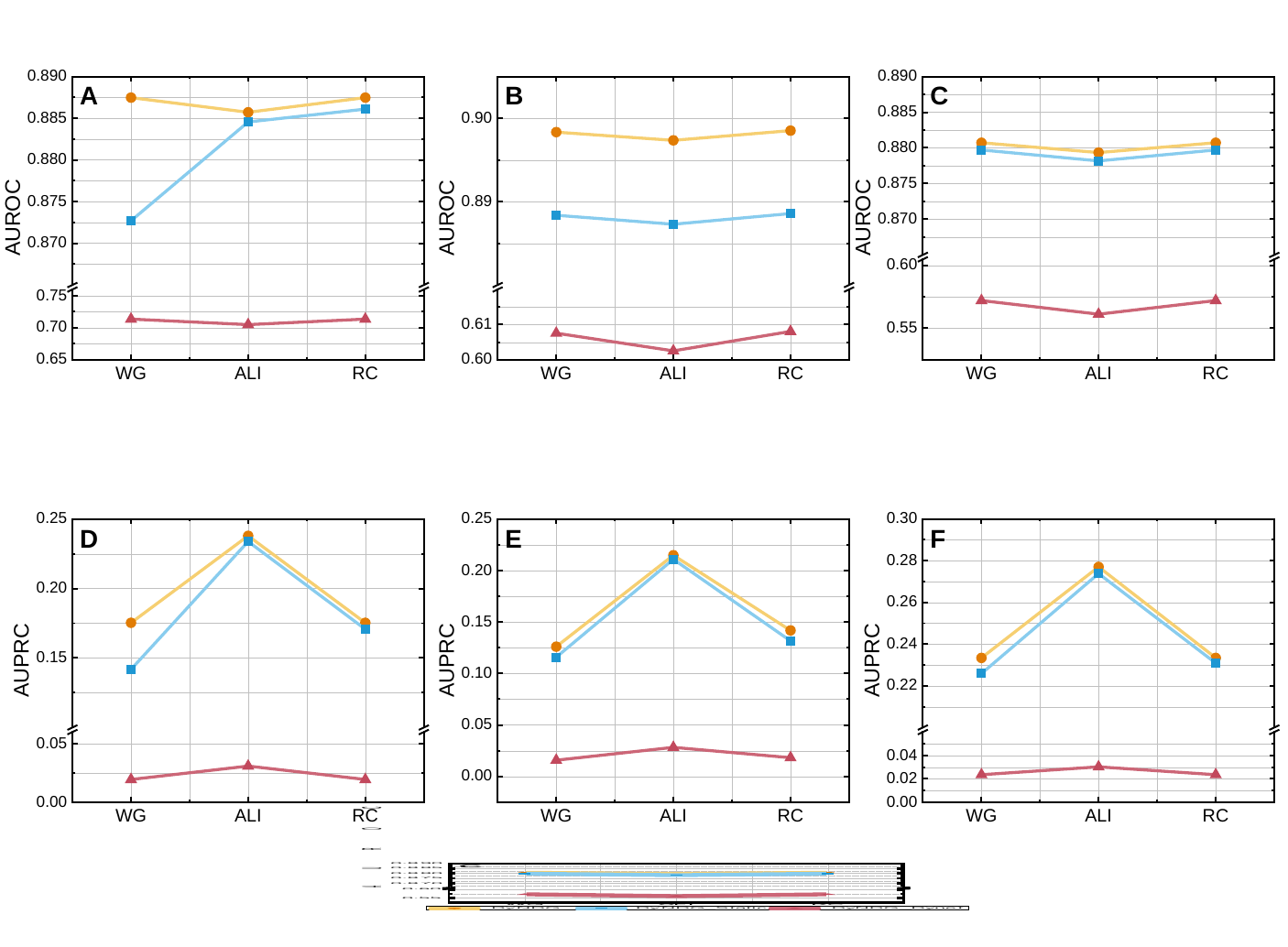

### Figure S16.jpg

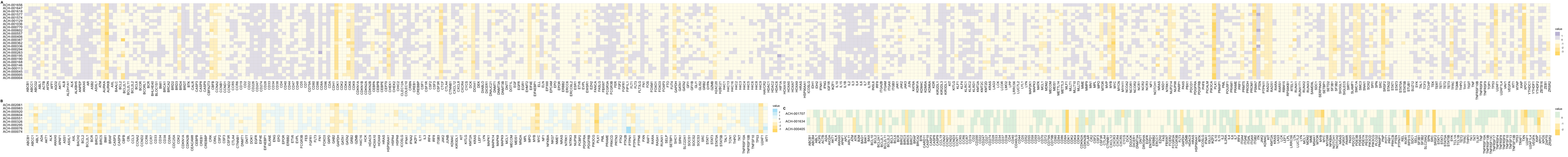
